# The insulin / IGF axis is critically important for controlling gene transcription in the podocyte

**DOI:** 10.1101/2024.05.20.594973

**Authors:** Jennifer A. Hurcombe, Lusyan Dayalan, Fern Barrington, Frédéric Burdet, Lan Ni, Joseph T. Coward, Mark Ibberson, Paul T. Brinkkoetter, Martin Holzenberger, Aaron Jeffries, Sebastian Oltean, Gavin I. Welsh, Richard J.M. Coward

## Abstract

Signalling to the podocyte via the structurally related insulin receptor (IR) or insulin-like growth factor 1 receptor (IGF1R) is important for podocyte function. This study sought to elucidate the compound role of the insulin/IGF1 axis in podocytes using transgenic mice and cell culture models deficient in both receptors.

Podocyte specific IR/IGF1R knockdown mice developed a severe kidney phenotype with albuminuria, glomerulosclerosis and renal failure with death occurring in some mice between 4 and 24 weeks. Simultaneous knockdown of both receptors in cultured podocytes resulted in >50% cell death by 7 days.

Proteomic analysis revealed a striking downregulation of spliceosome-related proteins in IR/IGF1R knockdown podocytes with long-read RNA sequence data indicating an increased fraction of transcripts with intron retention/premature termination codons in these cells. Furthermore, phospho-proteomic analysis revealed multiple insulin / IGF1 induced spliceosomal post-translational protein and kinase modifications suggesting dynamic control of this system.

This study underlines the importance of podocyte insulin/IGF signalling demonstrating a novel role for this extrinsic hormonal signalling axis in regulating gene transcription in this cell type.

## INTRODUCTION

Podocytes are highly specialised, terminally differentiated epithelial cells located at the urinary aspect of the glomerular basement membrane (GBM) in the kidney. They are crucial in maintaining glomerular filtration barrier integrity thus preventing the development of albuminuria, an independent risk factor for end-stage-kidney-disease [ESKD] ^1^, cardiovascular morbidity ^2^ and death. Therefore, understanding the critical signalling pathways necessary for maintaining podocyte health is desirable.

The insulin/insulin-like growth factor (IGF) signalling axis regulates many aspects of growth and metabolism: insulin signalling is predominantly associated with metabolism and IGF signalling with growth and cell division ^3^. However, insulin, IGF1 and their cognate receptors (the insulin receptor [IR] and insulin-like growth factor receptor [IGF1R]) share significant structural homology and several common downstream signalling nodes ^4^. The IR and IGF1R can bind each other’s ligands and can act redundantly in some contexts ^5,6^. The constitutive global (whole body) loss of either the IR or IGF1R in murine knockout models is lethal ^7,8^ suggesting a limited ability for these receptors to compensate for each other in every cell type. However, the insulin / IGF axis has different roles in different cell types. Several mouse models with tissue-specific dual knockout of both the IR and IGF1R show differential consequences occurring in a cell type dependent manner ^9–13^.

Insulin signalling to the podocyte via the IR is critical for podocyte function ^14–17^ and it is apparent that IGF signalling is also important for podocyte homeostasis ^18–20^. Recently we have shown that loss of podocyte IGF1 signalling can have both detrimental and beneficial effects dependent upon the level of IGF1R suppression: whilst near total loss of podocyte IGF1R expression results in severe mitochondrial dysfunction associated with cell death, partial receptor inhibition promotes podocyte survival in conditions of cellular stress ^21^.

Here we show that combined, simultaneous, loss of both the IR and IGF1R in podocytes is physiologically highly detrimental and present proteomic and transcriptomic analyses indicating that the IR/IGF1R hormonal signalling axis is crucial in controlling spliceosome assembly and RNA processing in this cell.

## METHODS

### Mouse models

Mice in which exon four of the IR ^7^ and exon three of the IGF1R have been flanked by loxP sites ^8^ were crossed with podCre mice ^22^. Progeny with the genotype podCre^+/0^;IR^wt/fl^;IGF1R^wt/fl^ was then crossed with IR^fl/fl^;IGF1R^fl/fl^ mice to generate podocyte specific IR/IGF1R knockdown animals from embryonic day 12 (podDKD) (Figure Supp 1A). Littermates served as controls. All mice were on a mixed genetic background including contributions from 129/Sv, FVB and C57BL/6. Both sexes were studied, and no phenotypic differences observed.

Transgenic mouse work was carried out in accordance with the University of Bristol’s institutional guidelines, and procedures approved by the United Kingdom (UK) Home Office in accordance with UK legislation. (Home Office Protocol numbers PPL 3003394 & PPL 2012285).

### Urinary albumin and creatinine measurements

Albumin and creatinine levels in spot collections of mouse urine were measured using a mouse-specific albumin ELISA (Universal Biologicals, Cat# E90-134) and creatinine companion kit (Exocell, Cat# 1012), following the manufacturer’s methodology.

### Histology and periodic acid-Schiff staining

Kidneys were fixed in 10% buffered neutral formalin, further processed and paraffin embedded. 3 μm sections were cut and stained using a periodic acid-Schiff staining kit (Sigma) according to the manufacturer’s instructions. Tissues were imaged using a Leica DN2000 microscope and micrographs taken using Leica Application Suite software. Image analysis was performed using ImageJ; all images were contrast enhanced using the same parameters.

Glomerulosclerosis was scored for each glomerulus as follows: 0=normal glomeruli; 1=up to 25% involvement; 2=25-50% involvement; 3=50-75% involvement; 4=over 75% involvement. The glomerulosclerosis index was calculated as described previously ^23^ and according to the formula [(1xN1)+(2xN2)+(3xN3)+(4xN4)]/(N0+N1+N2+N3+N4), where Nx is the number of glomeruli with each given score. Sections were assessed and scored blindly.

### Immunofluorescence

Frozen kidneys were sectioned at 5 μm. Sections were blocked in phosphate buffered saline (PBS) containing 3% bovine serum albumin (BSA) and 0.3% triton X-100 for 1 h, an antibody to Wilms Tumour Protein 1 (Abcam Cat#ab89901). Following 3 phosphate buffered saline (PBS) rinses, sections were incubated with a fluorophore-conjugated secondary antibody (Fisher) for 1 h at room temperature. Tissues were imaged using a Leica DM2000 microscope and micrographs taken with Leica Application Suite X software. Image analysis was performed with ImageJ; all images were contrast enhanced using the same parameters.

### Immunohistochemistry

Tissues were fixed in 10% buffered neutral formalin (Merck, Cat# HT501128), processed and paraffin embedded. 3 μm sections were deparaffinised in Histo-Clear II (National Diagnostics, Cat# HS-202) and rehydrated through a graded alcohol series. Antigen retrieval was in 10 mM citrate buffer, pH6 for by boiling for 10 min. Sections were quenched using 3% H2O2, followed by a blocking step in 3% normal goat serum for 30–45 min. Sections were incubated overnight at 4°C with an SF3B4 antibody (Novus Biologicals Cat#NBP-9269255) diluted 1:100. Sections were washed then incubated with SignalStain Boost detection reagent (Cell Signaling Technology, Cat# 8114) for 30 min at room temperature. SignalStain DAB substrate kit (Cell Signaling Technology, Cat# 8059) was applied for 1–2 min and the sections dehydrated and mounted in DPX (Sigma, Cat# 06522). Tissues were imaged using a Leica DM2000 microscope and micrographs taken with Leica Application Suite X software (Leica Microsystems). Image analysis was performed with ImageJ; all images were contrast enhanced using the same parameters.

### Electron microscopy

Tissues for electron microscopy were fixed in 0.1 M sodium cacodylate, 2% glutaraldehyde, and imaged on a Technai 12 transmission electron microscope. Average slit diaphragm, foot process and glomerular basement membrane width were calculated using ImageJ assessing at least 20 regions of glomerular basement membrane from at least 2 glomeruli per mouse.

### Lentiviral transduction of conditionally immortalised IR and/or IGF1R floxed podocyte cell lines

Kidneys were isolated from IR^fl/fl^;IGF1R^fl/fl^, IR^fl/fl^ and IGF1R^fl/fl^ mice and used to make homozygous floxed temperature-sensitive SV40 conditionally immortalised podocyte cell lines: double receptor knockdown (ciDKD), insulin receptor knockdown (ciIRKD) and IGF1 receptor knockdown (ciIGF1RKD) cells (Figure Supp 2A), using techniques as described previously ^24^. Podocytes were cultured at 33°C and when 50% confluent were thermo-switched to 37°C and incubated for a further 7 days before transduction with a lentivirus expressing Cre recombinase as described previously ^21^. Transduction was in RPMI media with hexadimethrine bromide (Sigma) at 4 μg/ml and the virus used at a multiplicity of infection of 1. Following a 24-hour incubation, the lentivirus was removed and replaced with fresh media. Cells were incubated for a further 3-7 days before imaging and protein extraction.

To determine cell number, cells were washed 3 times in PBS, the nuclei stained with Hoechst (Sigma) at 1 μg/ml and imaged using an IN Cell analyser 2200 (GE Healthcare), and data analysed using IN Cell analyser Developer software (GE Healthcare).

### Cell culture

Conditionally immortalised podocyte cell lines were cultured as described previously ^25^. For acute insulin and IGF1 stimulation, cells were serum starved for 4 hours, then 10 nM and 100 nM of insulin or 10 ng/ml and 100 ng/ml IGF1 was applied to the cells for 10 minutes. Spliceosome inhibitor treatment was carried out using pladienolide B at 0-100 nM for 48 hours before determination of cell number.

### Western blotting

Cultured cells were lysed in radioimmunoprecipitation assay buffer supplemented with protease and phosphatase inhibitors (Sigma). 10-30 μg of protein was resolved by electrophoresis and then transferred to a polyvinylidene difluoride membrane (Millipore). Membranes were blocked in TRIS-buffered saline with 0.1% Tween 20 and 5% BSA for 1 hour and then incubated overnight with primary antibody at a dilution of 1:1000. Primary antibodies used were from Cell Signalling Technology unless otherwise stated: IRβ Cat# 3025; IGF1R Cat# 9750; pAKT Cat#4060; AKT Cat# 2920; P-p44/42 MAPK Cat#4370; P44/42 MAPK Cat# 9102; EIF42A (Proteintech Cat#17564-1-AP); SF3B4 (Novus Biologicals cat#NBP-9269255); PTBP2 (Proteintech Cat#55196-1-AP). Membranes were washed before incubation with horseradish peroxidase conjugated secondary antibody (Sigma). Immunoreactive bands were visualised using Clarity ECL Western blotting substrate (Biorad) on a GE AI600 imager. Densitometry was performed using ImageJ software.

### Proteomic analysis

ciDKD, ciIRKD and ciIGF1RKD podocytes (3 days after Cre lentiviral transduction and before major cell loss)(Figure Supp 2B) were lysed in RIPA buffer and subjected to LC-MS/MS using isobaric TMT labelling as described previously ^26,27^. n=9 knockdown and control podocytes were analysed. Wild-type mouse podocytes served as controls.

The raw data files for the Total proteome analysis were processed and quantified using Proteome Discoverer software v2.1 (Thermo Scientific) and searched against the UniProt Mouse database (downloaded October 2019: 83152 entries) using the SEQUEST algorithm. Peptide precursor mass tolerance was set at 10 ppm, and MS/MS tolerance was set at 0.6 Da. Search criteria included oxidation of methionine (+15.9949) as a variable modification and carbamidomethylation of cysteine (+57.0214) and the addition of the TMT mass tag (+229.163) to peptide N-termini and lysine as fixed modifications. The data output from Proteome Discoverer 2.1 was further analysed in Microsoft Office Excel, Graphpad Prism 9 and Perseus 1.6.10.43 using log2-transformed scaled total protein abundance data. A *t*-test was used to determine significantly differentially expressed proteins. Hierarchical clustering was performed in Perseus by Euclidean distance using *k*-means pre-processing. Gene ontology (GO) and Kyoto Encyclopaedia of Genes and Genomes (KEGG) and enrichments were performed in Perseus using a Fisher test of significantly changed proteins versus the unchanged population after annotation with the proteins.

For Search Tool for the Retrieval of Interacting Genes/Proteins (STRING) enrichment analysis, the ciDKD expression dataset was submitted to STRING with log2 fold change values associated with each protein. Functional enrichments were identified, and enrichment scores calculated based on aggregate fold change or Kolmogorov-Smirnov testing in STRING.

### Phosphoproteomic analysis

ciDKD, ciIRKD, ciIGF1RKD cells were generated as described above. The phosphoproteome was assessed 3 days after initiating receptor knockdown with lentiviral Cre. Wild-type (receptor sufficient conditionally immortalised murine podocytes) were also studied in the same manner.

Cells were starved of insulin and IGF1 for 4 hours and then stimulated with either 10 nM Insulin or 10 ng/ml IGF1 for 10 minutes. The raw data files for the phosphoproteome analyses were processed and quantified using Proteome Discoverer software v2.1 (Thermo Scientific) and searched against the UniProt Mouse database using the SEQUEST algorithm. Peptide precursor mass tolerance was set at 10ppm, and MS/MS tolerance was set at 0.6 Da. Search criteria included oxidation of methionine (+15.9949) as a variable modification and carbamidomethylation of cysteine (+57.0214) and the addition of the TMT mass tag (+229.163) to peptide N-termini and lysine as fixed modifications. For the phosphoproteome analysis, phosphorylation of serine, threonine and tyrosine (+79.966) was also included as a variable modification. Searches were performed with full tryptic digestion and a maximum of 2 missed cleavages were allowed. The reverse database search option was enabled and all data was filtered to satisfy false discovery rate (FDR) of 5%. The data output from the Proteome Discoverer 2.1 analysis was processed in the following ways. The annotation of master proteins was improved using an in-house R script which integrates Uniprot Annotation levels into the selection of master proteins. Scaled Protein abundance data were log2 transformed to bring them closer to a normal distribution. Phosphopeptide abundance data were calculated by summing the abundances of identical phosphorylated PSM sequences into a single Phosphopeptide abundance value. The Proteome Discoverer software normalises the data for the Total Proteome analysis such that the total peptide abundance for each sample is the same. These normalisation factors were then applied to the raw phosphopeptide abundances for each sample (generated above). Each phosphopeptide was then further normalised to the observed abundance of its corresponding protein, such that any changes observed reflected changes in protein phosphorylation rather than in protein abundance. To identify differentially expressed proteins and phosphopeptides, a paired *t*-test was conducted between conditions of interest with a cut-off of p<0.05.

### Long-read RNA sequencing

RNA was isolated from ciDKD podocytes (3 days after Cre lentiviral transduction) and wild-type control cells (n=4 for each group) using an RNeasy mini kit (Qiagen). RNA was quantified using a Qubit Fluorimeter (Thermo Fisher Scientific) and quality assessed using the TapeStation system (Agilent). All samples had an RNA integrity number >8.

Barcoded PCR-cDNA library preparation was performed on 75 ng of RNA from each sample using the SQK-PCB109 kit (Oxford Nanopore Technologies). The resulting library was quantified using a Qubit Fluorimeter (Thermo Fisher Scientific) and molecular weight estimated using the TapeStation system (Agilent). 50 fmol of library was then loaded onto three promethION R9.4.1 flowcells (Oxford Nanopore Technologies), sequenced for 72 h and basecalled using guppy_basecaller 5.0.17 (Oxford Nanopore Technologies) in high accuracy mode.

### Long-read RNA analysis

IsoQuant analysis: the reads were aligned using IsoQuant (version 2.3.0), with option --data_type nanopore. The mouse gencode annotation (V29) ^28^ was used. The IsoQuant reference-based transcripts counts were used for further analysis. Differential expression analysis between the 2 conditions was performed using limma function with voom approach from limma Bioconductor package ^29^. The transcripts were analyzed without being grouped by genes, which will lead to a more stringent analysis than if done by gene (multiple correction testing is done by transcript). For visualization in the genome browser, the bam files were converted to bigWig files (using a scaleFactor calculated with edgeR ^30^). FLAIR analysis (flair version 1.6.4): pychopper (version 2.7.2) ^31^ was first run on the total reads. Then the full length and rescued reads were aligned with “flair align” on the reference fasta file. The alignments were corrected using “flair correct”. The alignments were then collapsed using “flair collapse”, using the 8 replicates, into 1 complete reference. Then flair quantify, diffExp and diffSplice were run using the 2×4 replicates to get the rank lists of the 4 splicing event types (cassette, 3’, 5’, IR). A custom R script was used to find the nearest gene to the splicing events. In parallel, flair mark_intron_retention and predictProductivity (option --longestORF) were run on each corrected alignments to list the intron retention events by sample and predict the transcripts productivity by sample.

### Quantitative PCR

cDNA was synthesised from RNA isolated from ciDKD and wild-type podocytes using the High-Capacity RNA-to-cDNA kit (ThermoFisher Scientific). Quantitative PCR was performed using SYBR Green JumpStart Taq ReadyMix (Merck) in a StepOnePlus PCR machine (Applied Biosystems). Primers with the following sequences were purchased from Life Technologies: Fn1 F 5’ CCCAGCTCACTGACCTAAGC 3’; Fn1 R 5’ GGAAGAGTTTAGCGGGGTCC 3’; Hcfc1r1 F 5’ GCCACCACTGGGGTAACTC 3’; Hcfc1r1 R 5’CTTCGGGAAAAGTCACAGGG3’; beta actin F 5’ GACAGGATGCAGAAGGAGATTACT 3’; beta actin R 5’ TGATCCACATCTGCTGGAAGGT 3’.

### Statistical analysis

Statistical analysis was performed in Graphpad Prism 9 and data presented as the mean +/− SD unless otherwise stated. Statistical significance was calculated with *t*-tests to compare two groups or one-way ANOVA to compare more than two groups and taken as p<0.05.

### Data availability

The authors declare that all data supporting the findings of this study are available within the article and its supplementary information files or from the corresponding author upon reasonable request. The mass spectrometry proteomics data have been deposited in the PRIDE repository as part of the ProteomeXchange Consortium^32^ under the dataset identifier PXD051018.

## RESULTS

### podDKD mice develop a severe renal phenotype by 24 weeks

To study the effect of the combined loss of podocyte IR and IGF1R *in vivo*, transgenic mice were generated by crossing IR^fl/fl^;IGF1R^fl/fl^ mice with mice expressing Cre recombinase under the control of a podocin promotor^22^ (Figure Supp 1A). There was no difference in body weight or blood glucose in podDKD mice compared with littermate controls (Figures 1A and B). However, knockdown mice developed albuminuria with significantly increased urinary albumin:creatinine ratio (uACR) (p<0.01) at 24 weeks of age (Figure 1C and Figure Supp 1B). The severity of the phenotype was variable and while some mice exhibited only mild albuminuria, more severe renal disease was observed in others resulting in death in ∼20% of the mice. Of note, we found variable levels of receptor knockdown in the models which we suspect contributed to the variable phenotype observed (Figure Supp 1C). We considered using an inducible podocyte specific cre driver to eliminate any developmental confounders associated with the constitutive podocin cre^22^ model, but the level of floxed gene excision is less in these systems compared to constitutive cre driven models (Figure Supp 1D), so we concluded that the podocyte IR and IGF1R levels would be even less “knocked-down” using this approach. uACR was not increased in Cre negative and podCre expressing mice at 6 months excluding Cre toxicity as the driver of albuminuria in podDKD mice (Figure Supp 1E). Glomerulosclerosis was significantly increased (p<0.05) in podDKD mice with areas of sclerosis and tubular protein casts visible on periodic acid-Schiff (PAS) stained sections (Figure 1D), together with evidence of increased fibrosis as determined by Masson’s trichrome staining (Figure 1E).Transmission electron microscopy (TEM) analysis revealed disruption of the filtration barrier ultrastructure with significantly increased (p<0.01) foot process width (Figure 1F). A limited number of serum creatinine levels were recorded at 4-6 months and showed no significant difference (Figure Supp 1F). Podocyte number, evaluated by immunofluorescence (IF) using a podocyte specific WT1 antibody, was significantly (p<0.0001) reduced in podDKD mice at 6-months (Figure 1G).

**Figure 1.**
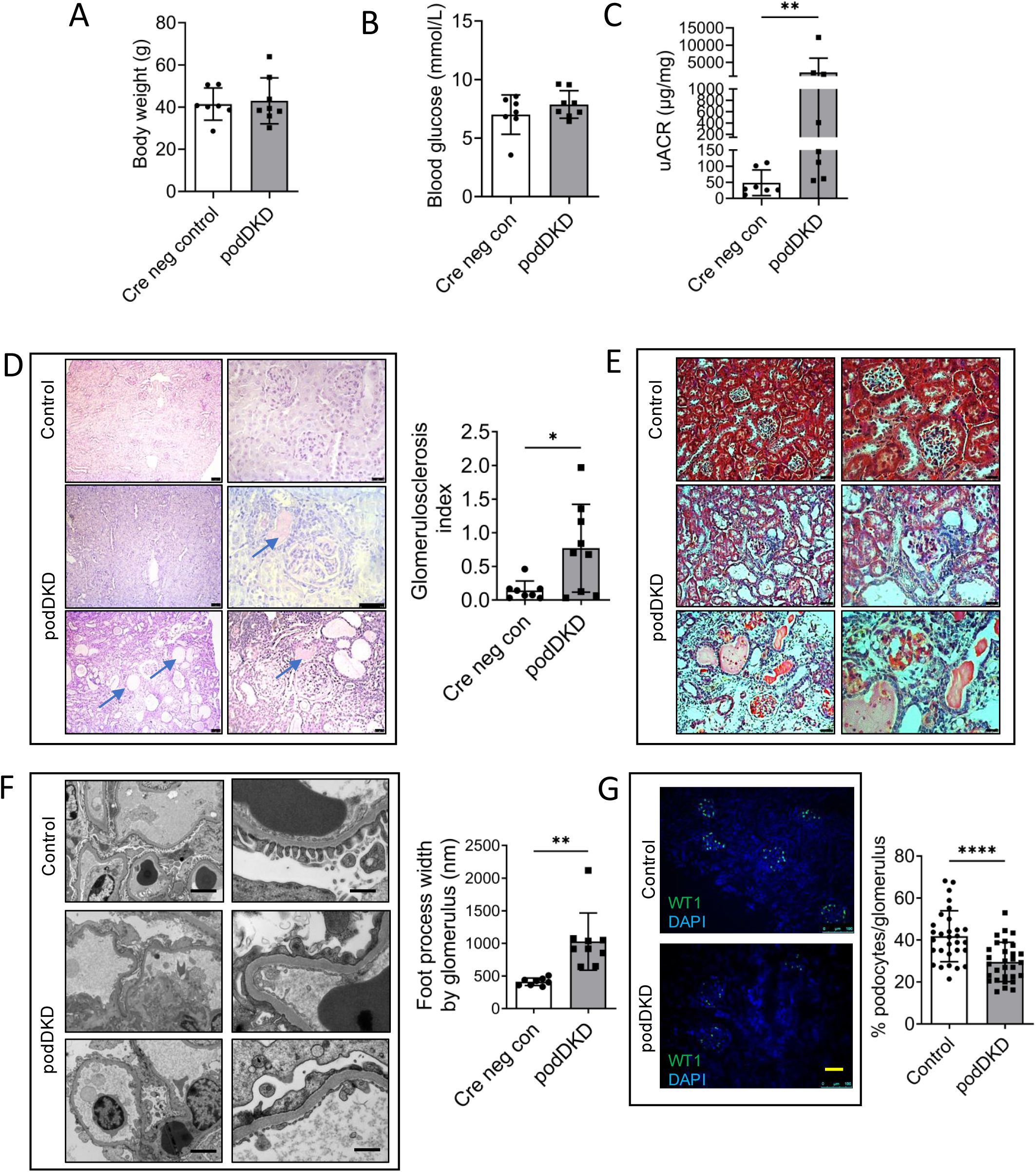
podDKD mice develop a renal phenotype by 24 weeks. **A.** and **B**. Body weight (**A**) and blood glucose (**B**) are not significantly different in podDKD mice at 24 weeks compared with littermate controls (n=7-8 each group). **C.** Urinary Albumin Creatinine ratio (uACR) is significantly increased in podDKD mice at 24 weeks. Unpaired *t*-test, **p<0.01 (n=7-8 mice per group). **D.** Images and quantification of PAS staining show tubular protein casts (indicated by arrows) and glomerulosclerosis in podDKD mice. Scale bar=25 μm. Unpaired *t*-test *p<0.05. **E.** Masson’s trichrome staining shows increased fibrosis (blue staining) in podDKD mice at 24 weeks. Scale bar=25 μm. **F.** Transmission Electron Micrograph (TEM) images of glomerular filtration barrier (GFB) (scale bar left panels=5 μm, right panels =500 nm) show ultrastructural damage to the GFB in podDKD mice with significantly increased foot process width. Unpaired *t*-test **p<0.01. **G.** Immunofluorescent staining and quantification of WT1 in podDKD and Cre negative control mice at 24 weeks of age shows significant reduction in % podocytes per glomerulus in podDKD mice. Nuclei counterstained with DAPI. Scale bar=100 μm. Unpaired *t*-test ****p<0.0001, ≥8 glomeruli analysed per mouse, 3 mice per group. Scale bar=100 μm.

### Simultaneous knockdown of podocyte IR and IGF1R *in vitro* is highly detrimental

We surmised that the variable phenotype observed in our transgenic mouse model was predominantly due to differences in the efficiency of Cre mediated excision and level of receptor knockdown. With the aim of developing a model with near total loss of the IR and IGF1R to use in *in vitro* studies, a conditionally immortalised podocyte cell line was generated from IR^fl/fl^;IGF1R^fl/fl^ mice and these cells subsequently transduced with Cre expressing lentivirus (ciDKD) (Figure Supp 2A). This allowed us to initially culture the cells containing both receptors before their simultaneous knock-down using lentiviral delivered Cre recombinase. Western blot analysis showed a highly significant decrease of the IR (p<0.0001) and IGF1R (p<0.0001) in ciDKD podocytes of >80% (Figure 2A). Phosphorylation of AKT and p44/42 MAPK in response to either acute insulin or IGF1 stimulation was reduced (p<0.05) indicating suppression of insulin/IGF1 signalling in ciDKD podocytes (Figures 2B and C). Compound loss of both the IR and IGF1R was detrimental, resulting in >50% podocyte loss (p<0.0001) 7 days after Cre-mediated gene excision (Figure 2D).

**Figure 2.**
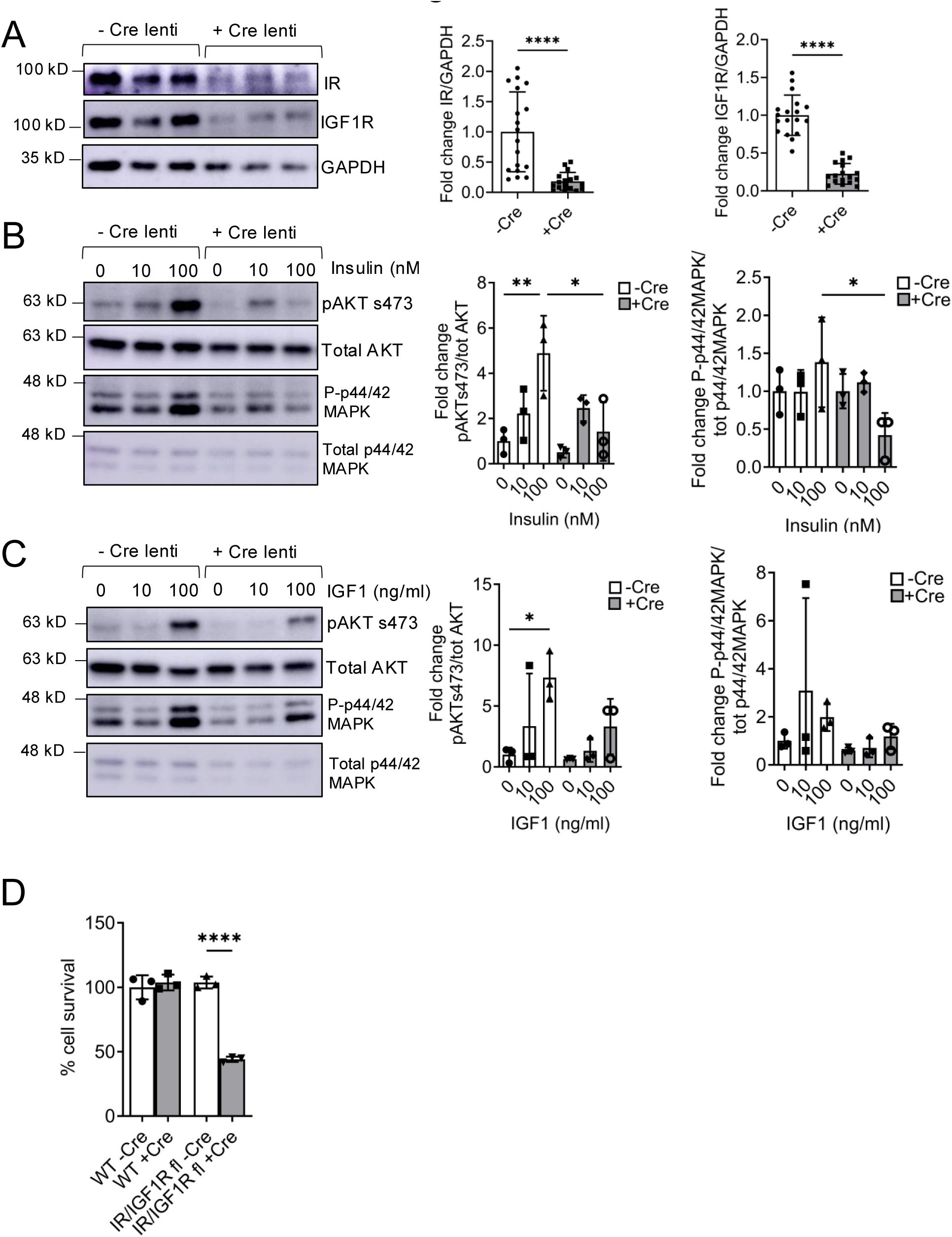
Simultaneous knockout of podocyte IR and IGF1R *in vitro* is highly detrimental. **A.** Western blot shows >80% reduction of IR and IGF1R protein in ciDKD cells. *t*-test, ****p<0.00001, n=18. **B.** Phosphorylation of AKT and p44/42MAPK in response to acute insulin stimulation at 10 nM and 100 nM for 10 minutes was significantly reduced in ciDKD podocytes. One way ANOVA, **p<0.01, *p<0.05, n=3. **C.** Phosphorylation of AKT and p44/42MAPK in response to acute IGF1 stimulation at 10 and 100 ng/ml for 10 minutes was reduced in ciDKD podocytes. One way ANOVA, *p<0.05, n=3. **D.** Fewer than 50% of ciDKD cells survive 7 days after gene excision. *t*-test, ****p<0.0001, n=3-4 independent experiments.

### Proteomic analysis of ciDKD podocytes reveals downregulation of spliceosome proteins

To elucidate the mechanistic processes regulated by insulin/IGF1 signalling in podocytes, we performed unbiased tandem mass tagged (TMT) LC MS/MS proteomics to compare total protein expression in ciDKD compared with wild-type cells (Figure 3A). We performed this 3-days after Cre exposure when cell death was approximately 20% compared to the 50% cell death observed at 7 days (Figure S2B). Hierarchical clustering and principal component analysis (PCA) was performed using Perseus software to determine that samples were clustered according to the experimental group (Figures Supp 3A and Supp 3B). 4842 proteins were significantly differentially expressed in ciDKD cells compared to wild-type controls with 2127 proteins upregulated and 2715 proteins downregulated (Figure 3A and Figure Supp 3C). Perseus software was used to perform hierarchical clustering and identify Gene Ontology (GO) and Kyoto Encyclopaedia of Genes and Genomes (KEGG) terms enriched in ciDKD podocytes (Figure 3B). This revealed a downregulation of proteins involved in DNA replication and cell cycling along with a striking downregulation of proteins involved in spliceosome function and RNA processing (Figure 3B). These findings were confirmed using Search Tool for the Retrieval of Interacting Genes/Proteins (STRING) to identify KEGG terms overrepresented in the ciDKD proteome (Figure 3C). STRING identified the spliceosome as the most significantly enriched KEGG term in dual receptor knockout cells with a log10 false discovery rate ∼2 fold greater than any other (Figure 3D).

**Figure 3.**
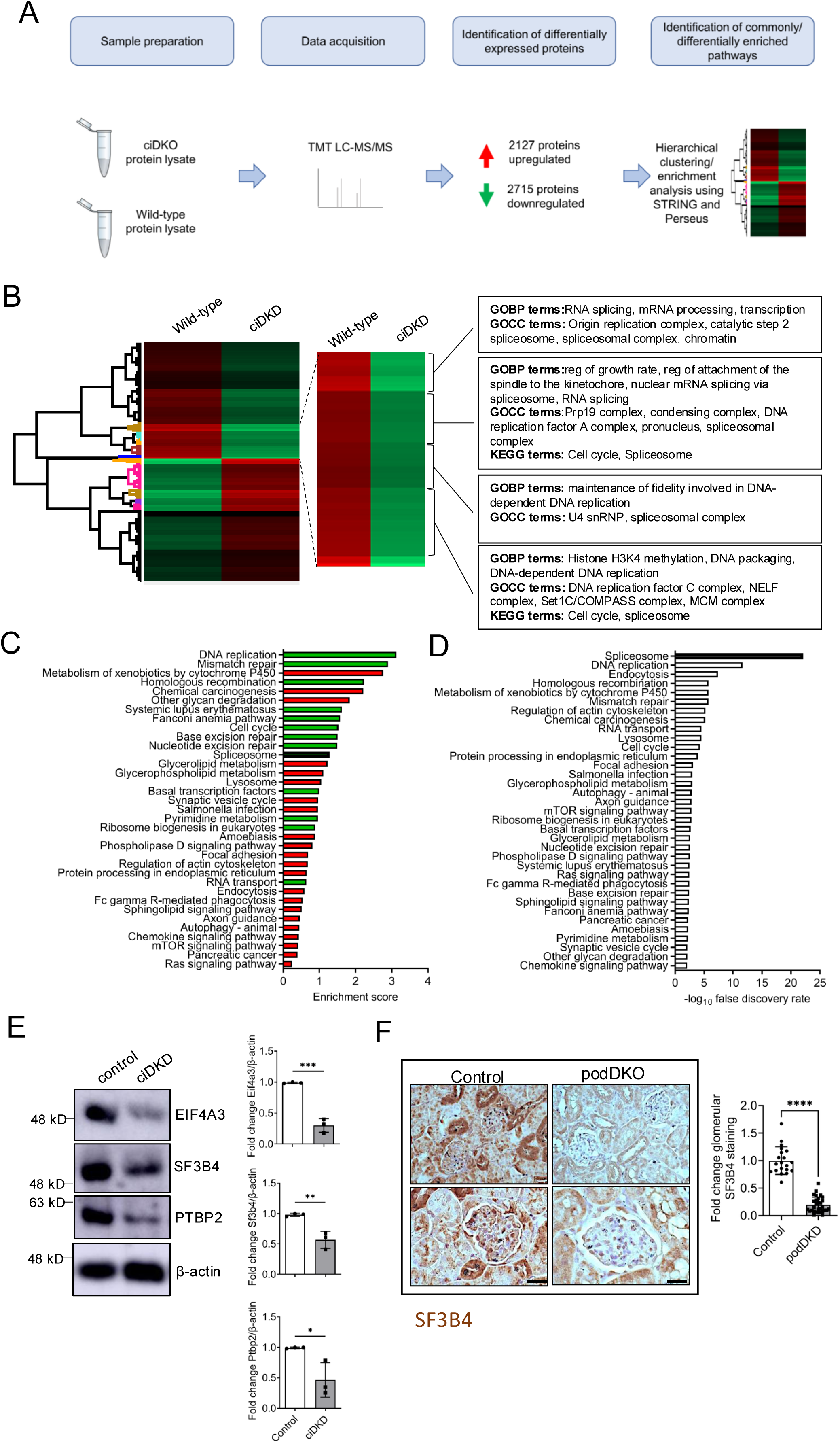
Proteomic analysis of ciDKD podocytes reveals downregulation of spliceosomal proteins. **A.** Schematic outlining workflow for proteomic analysis. **B.** Heat map showing hierarchical clustering of ciDKD vs wild-type podocyte proteomes (decreased protein expression in green, increased expression in red) and the GO and KEGG terms enriched in four major clusters. **C.** and **D.** KEGG enriched terms in STRING. Downregulated pathways (green) are associated with higher enrichment scores in comparison to upregulated pathways (red). Enrichment scores are computed by STRING using the Kolmogorov-Smirnoff test. KEGG term “Spliceosome” (in black) is associated with a high enrichment score and the highest false discovery rate across all terms. **E.** Western blots show significantly reduced levels of EIF4A, SF3B4 and PTBP2 in ciDKD podocytes compared with wild-type cells. Unpaired *t*-test, ***p<0.001, **p<0.01, *p<0.05, n=3 independent experiments. **F.** Representative immunohistochemistry and quantification using an antibody to Sf3b4 shows reduced expression in the glomeruli of podDKD mice compared to littermate controls. *T*-test ****p<0.0001.

To validate results from the proteomic data, western blot analysis of wild-type and ciDKD podocytes was performed using antibodies to selected spliceosome proteins. This confirmed that levels of eukaryotic initiation factor-4A (EIF4A) (p<0.001), splicing factor 3B subunit 4 (SF3B4) (p<0.01) and polypyrimidine tract binding protein 2 (PTBP2) (p<0.05) were significantly reduced in knockout cells (Figure 3E). We also validated proteomic changes occurring in our cell model were present in our animal model. Immunohistochemical glomerular SF3B4 expression was significantly lower in podDKD mice compared to littermate controls (Figure 3F).

### Exposure of cultured podocytes, but not glomerular endothelial cells, to the spliceosome inhibitor pladienolide B results in dose dependent cell death

Proteomic analysis indicated a significant reduction in spliceosome proteins associated with all stages of the splicing process in ciDKD podocytes (Figure 4A). We therefore hypothesised that impaired spliceosome assembly and function may be at least partially responsible for the deleterious phenotype observed with podocyte IR/IGF1R knockdown. To determine the importance of spliceosome activity to mature podocyte homeostasis and to ascertain whether different cell types are equally sensitive to spliceosome inhibition, the cytotoxic effect of pladienolide B (a spliceosome inhibitor targeting SF3B) ^33^ was compared in HeLa cells, fully differentiated, wild-type murine podocytes and glomerular endothelial cells. Consistent with previous observations ^33^, exposure of HeLa cells to pladienolide B at 0-100 nM for 48 hours was cytotoxic at ≥1 nM (Figure 4B).

**Figure 4.**
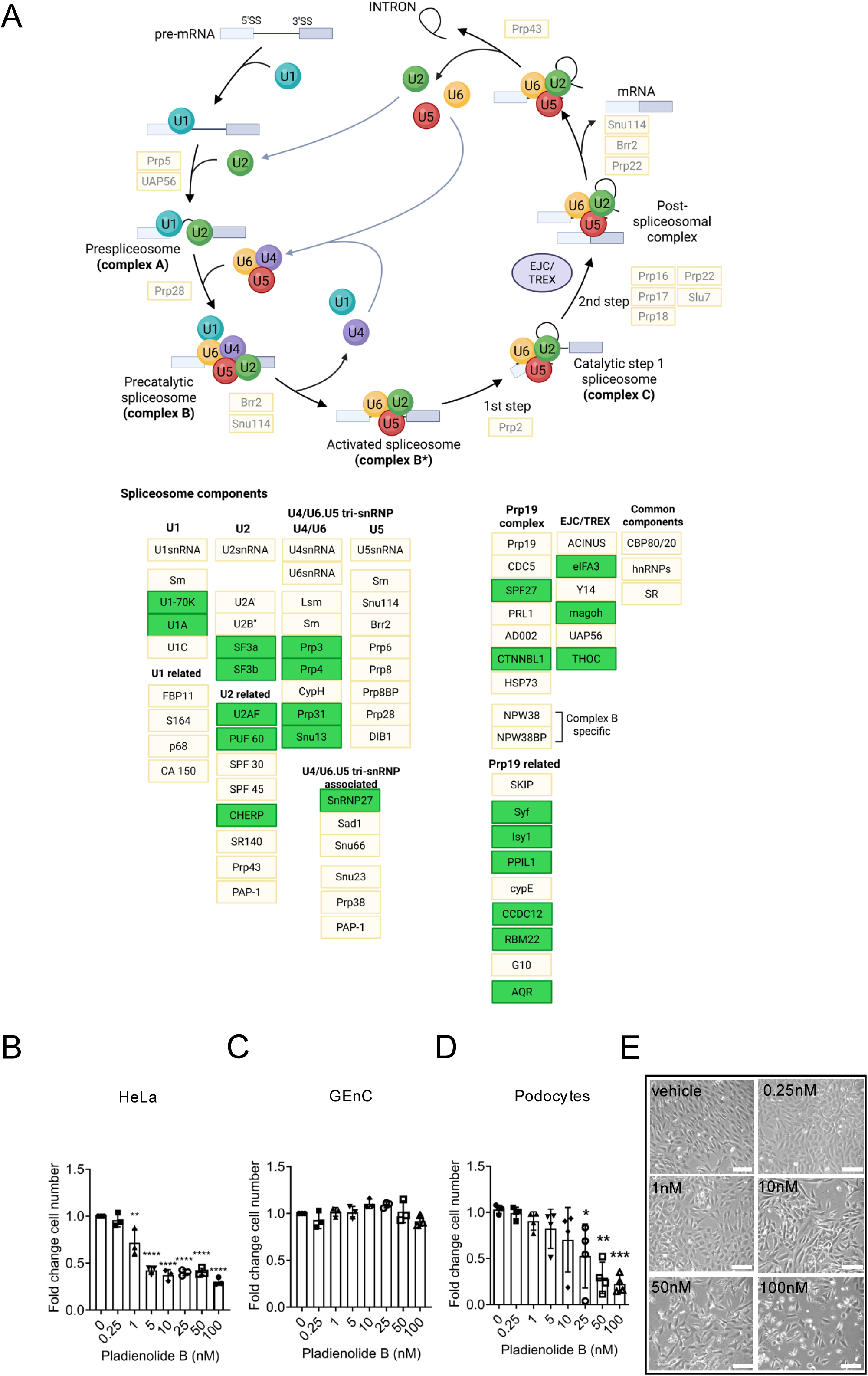
Exposure of cultured podocytes, but not glomerular endothelial cells, to the spliceosome inhibitor pladienolide B results in dose dependent cell death. **A.** Schematic overview of the spliceosome pathway showing a majority of spliceosome proteins are significantly downregulated (green) in ciDKD podocytes. **B-D.** Pladienolide B exposure for 48 hours in HeLa cells (**B**), glomerular endothelial cells (GenC) (**C**) and podocytes (**D**). One way ANOVA, ****p<0.0001, **p<0.01, *p<0.05, n=3 independent experiments. **E.** Bright field images of wild-type podocytes exposed to the indicated concentrations of pladienolide B. Scale bar=100 μm.

Pladienolide B-induced spliceosome inhibition also resulted in a significant (p<0.05) dose dependent loss of podocytes at a concentration of ≥25 nM (Figure 4D and E). In contrast, no cytotoxicity was observed in glomerular endothelial cells (Figure 4C). As previously stated, we also found decreased levels of SF3B4 in podocytes from podDKD mice (Figure 3F).

### Loss of podocyte IR and IGF1R is associated with increases in intron retention and unproductive transcript expression

To obtain an overview of global differences in various alternative splicing events in the ciDKD transcriptome, long-read RNA sequencing was used. Long-read sequencing was preferred as short-read RNA seq tools have been found to detect such events with poor accuracy ^34^. RNA was isolated from ciDKD and wild-type control cells, sequenced using long-read RNA Oxford Nanopore PromethION flow cells (Figure 5A and Figure Supp 4A-C) and the resulting data analysed using the Full-Length Alternative Isoform Analysis of RNA (FLAIR) tool. An overview of the main types of alternative splicing events is depicted in Figure 5B and rank lists of genes in ciDKD podocytes with significantly increased (adjusted p-value <0.05) intron retention splicing; alternative 5’ splicing; alternative 3’ splicing and cassette exon splicing are shown in Supp Tables 1-4. KEGG pathway enrichment analysis using STRING, shows an overlap in overrepresented pathways (including spliceosome, RNA transport and ribosome) from rank lists of genes with intron retention, cassette exon splicing and alternative 5’ splicing (Figure Supp 4D-F).

**Figure 5.**
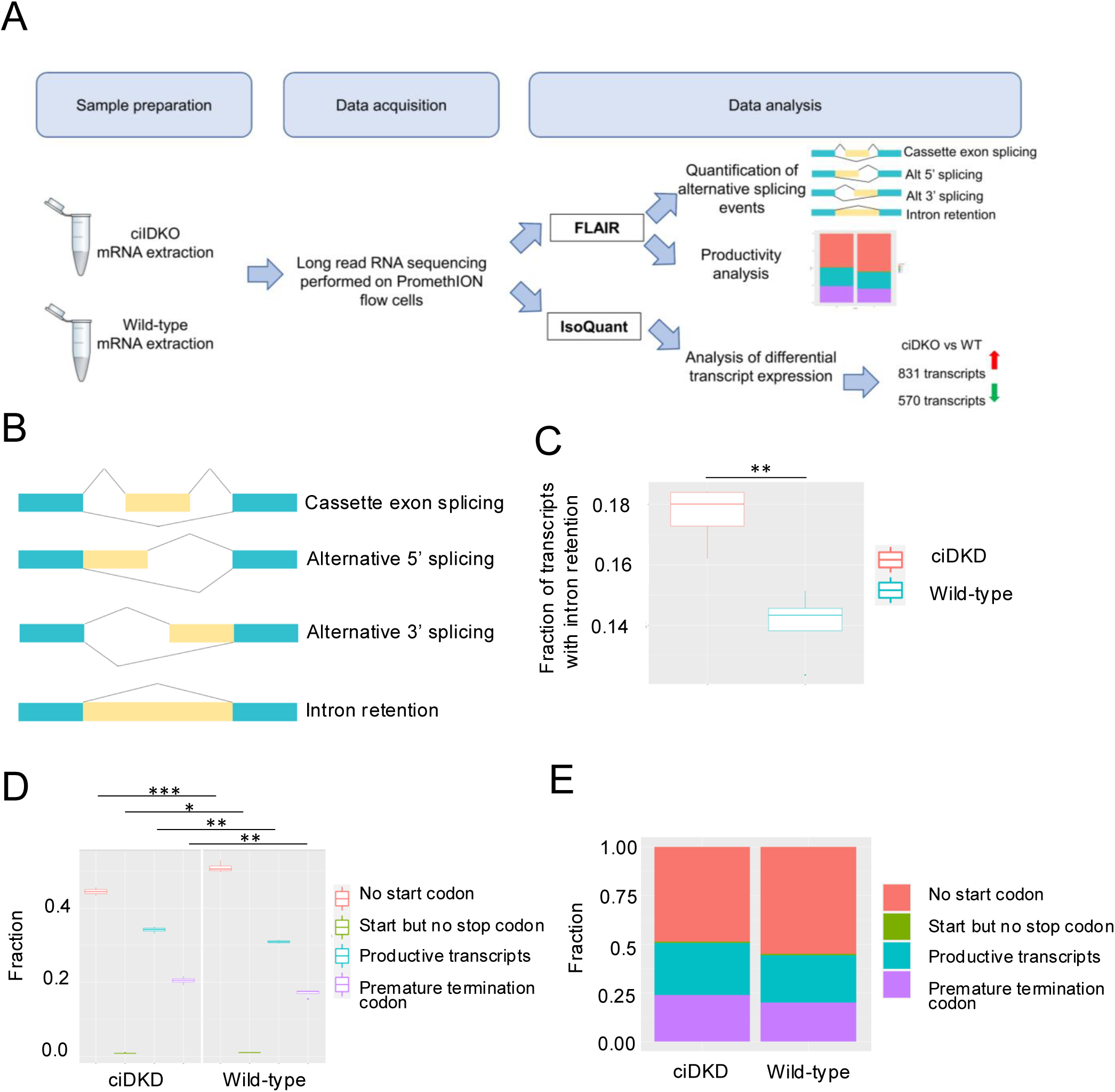
Loss of podocyte IR and IGF1R is associated with increases in intron retention and unproductive transcript expression. **A.** Schematic outlining long-read RNA sequencing workflow. **B.** An overview of the alternative splicing events quantified by FLAIR. Boxes represent exons (blue=constitutive exons; yellow=alternative exons), lines represent introns. **C.** Boxplot showing the fraction of transcripts with intron retention events in ciDKD and wild-type podocytes. *t*-test, **p<0.01, n=4 for each experimental condition. **D.** Box plot of the fraction of productivity events in the ciDKD and wild-type transcriptomes shows a higher proportion of unproductive transcripts in ciDKD podocytes. Includes no start codon; start codon but no stop codon; productive transcripts; premature termination codon i.e. unproductive transcript. *t*-test, ***p<0.001, **p<0.01, *p< 0.05, n=4 for each experimental condition. **E.** Proportional stacked bar graph of productivity events in the ciDKD and wild-type transcriptomes.

**TABLE 1.**

| Gene name | Transcript ID | Log FC | P. Value | adj.P.Val | Transcript type |
| --- | --- | --- | --- | --- | --- |
| D130062J10Rik | ENSMUST00000223763.2 | -3.8196361 | 0.0000000 | 0.0000085 | lncRNA |
| Fn1 | ENSMUST00000186879.2 | -6.9625749 | 0.0000000 | 0.0000010 | retained intron |
| Fn1 | ENSMUST00000055226.13 | 1.1097711 | 0.0060383 | 0.0509845 | protein coding |
| Fn1 | ENSMUST00000189821.7 | -0.8737398 | 0.0359924 | 0.1538286 | protein coding |
| Fn1 | ENSMUST00000185408.7 | 2.6214948 | 0.1620950 | 0.3828405 | retained intron |
| Fn1 | ENSMUST00000190780.7 | -0.4741740 | 0.2093201 | 0.4430398 | protein coding |
| Fn1 | ENSMUST00000187938.7 | 0.3681171 | 0.3633759 | 0.6025781 | protein coding |
| Fn1 | ENSMUST00000186129.7 | -0.1591273 | 0.5248095 | 0.7369268 | protein coding |
| Fn1 | ENSMUST00000189160.2 | 0.1149073 | 0.7523422 | 0.8668309 | retained intron |
| Fn1 | ENSMUST00000188894.7 | 0.0545639 | 0.8697479 | 0.9320835 | protein coding |
| Gm13196 | ENSMUST00000118297.2 | 7.0682661 | 0.0000000 | 0.0000043 | processed<br>pseudogene |
| Gm43461 | ENSMUST00000202351.2 | -2.5532991 | 0.0000000 | 0.0000127 | TEC |
| Gsta4 | ENSMUST00000034903.7 | 2.8474040 | 0.0000000 | 0.0000079 | protein coding |
| Gsta4 | ENSMUST00000213215.2 | 0.2810244 | 0.6215751 | 0.7841542 | protein coding |
| Hcfc1r1 | ENSMUST00000179928.2 | -7.5317703 | 0.0000000 | 0.0000079 | protein coding |
| Hcfc1r1 | ENSMUST00000180140.8 | 1.7727865 | 0.0098495 | 0.0692495 | protein coding |
| Hcfc1r1 | ENSMUST00000024697.5 | 0.4134073 | 0.0171869 | 0.0973040 | protein coding |
| Id3 | ENSMUST00000008016.3 | 2.6838864 | 0.0000000 | 0.0000079 | protein coding |
| Lamtor3-ps | ENSMUST00000164408.8 | 6.7070388 | 0.0000000 | 0.0000098 | protein coding |
| Marcks | ENSMUST000000092584.6 | 5.8763865 | 0.0000000 | 0.0000001 | protein coding |
| Rpl15-ps2 | ENSMUST000000184247.2 | 13.8135821 | 0.0000000 | 0.0000001 | processed<br>pseudogene |

To compare the intron retention in the ciDKD versus controls, the number of intron retention events by sample were counted using FLAIR mark_intron_retention, and then divided by the number of transcripts per sample. The ciDKD samples had a significantly (p-value = 0.004) higher proportion of transcripts with intron retention: ∼18% of transcripts in ciDKD podocytes compared with 14% in wild-type controls (Figure 5C). The overlap of the intron retention events between experimental repeats is ∼30% in both wild-type and ciDKD podocytes. The transcript productivity was also assessed using FLAIR predictProductivity with transcripts classified in 4 categories: PRO (productive); PTC (premature termination codon, i.e. unproductive); NGO (no start codon) and NST (has start codon but no stop codon appended to the end of the isoform name). The fraction of transcripts harbouring a premature termination codon is significantly higher (p<0.001) in ciDKD podocytes (Figure 5D and E).

### Compound knockdown of the IR and IGF1R alters the podocyte transcriptome

To determine the effect of spliceosome depletion on the transcriptomic profile of ciDKD podocytes, IsoQuant ^35^ was used to quantify genes and transcripts from existing GENCODE mouse annotation, then limma ^29^ used to determine differential gene and transcript expression. The number of reads per sample is shown in Figure Supp 4A. Variance-stabilized transformed RNA Seq data (Figure Supp 4B) were used in principal component analysis and confirmed that samples were clustered according to the experimental group (Figure Supp 4C). The number of minimally expressed transcripts in the experiment was 31580 and using an adjusted p-value threshold of 0.01, 831 transcripts were significantly upregulated and 570 significantly downregulated in ciDKD podocytes. The top 10 most significantly differentially expressed genes (analysed by transcript) are shown in Table 1. These included down-regulation of host cell factor C1 regulator 1 (Hcfc1r1), a nuclear export factor which regulates the activity of the transcription factor host cell factor 1 (Hcf1) and which analysis using FLAIR identified as subject to increased alternative 3’ splicing in ciDKD podocytes (Supp Table 3). Also, fibronectin 1 (Fn1) encoding an extracellular matrix protein which exists as different isoforms derived by alternative splicing at three sites: Extra Domains A and B exons (EDA and EDB) and the type III connecting segment ^36^. Fn1 splice variants were differentially expressed in ciDKD podocytes with increased expression (log fold change of 1.11[=2.16 fold]) of an Fn1 transcript containing the EDA and EDB exons (Figure 6A-C). qPCR validation confirmed the decrease in Hcfc1r1 expression (Figure 6D) while primers designed to specifically amplify transcripts containing the detrimental fibrosis-associated Fn1 EDB exon also verified that expression of such transcripts is significantly increased (1.8 fold, p<0.001) in ciDKD podocytes when compared with wild-type control cells (Figure 6E).

**Figure 6.**
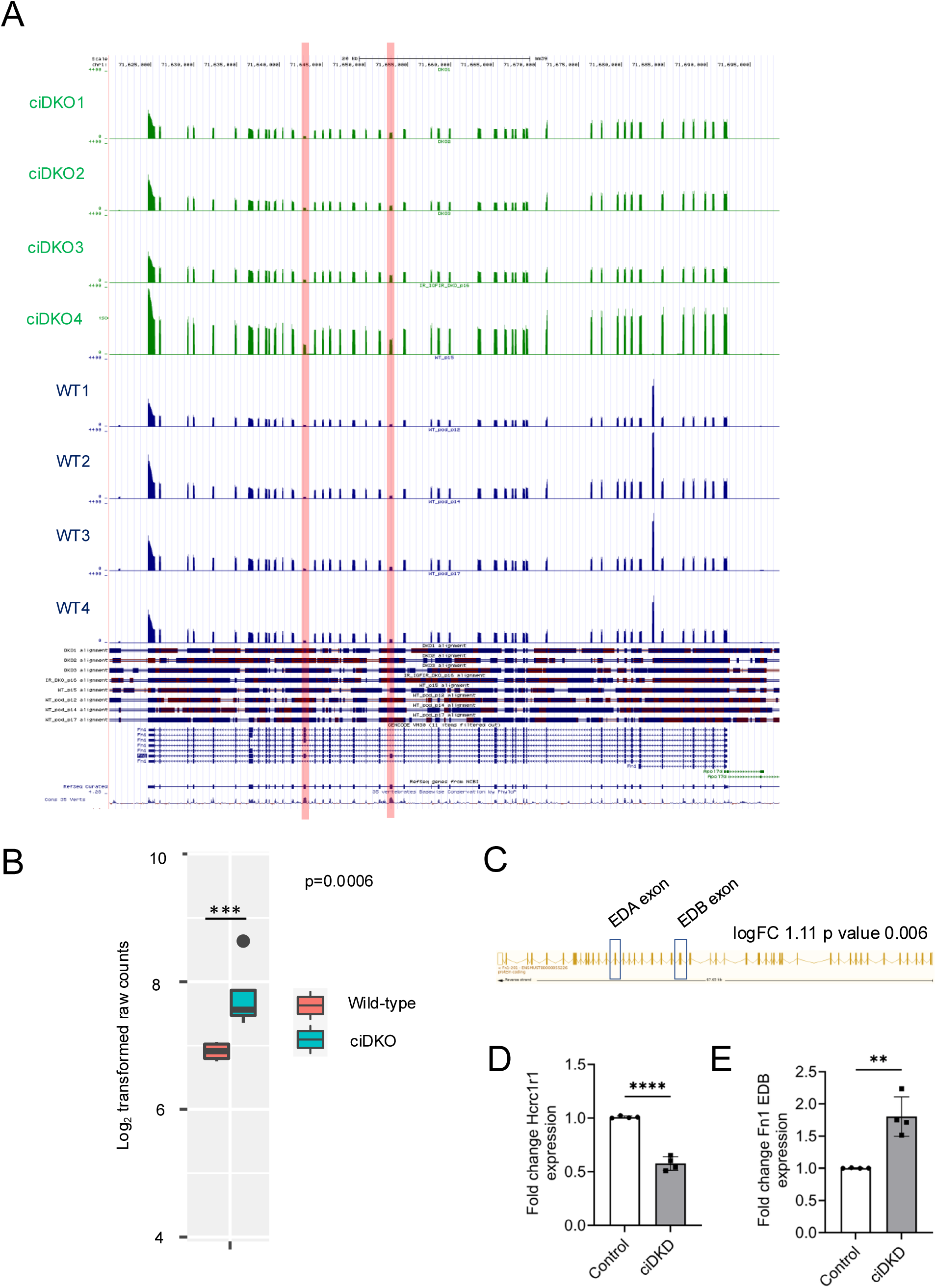
Compound knockdown of the IR and IGF1R alters the podocyte transcriptome. **A.** UCSC genome browser tracks of transcripts annotated to *Fn1.* Red lines indicate the number of reads in each sample containing the EDA and EDB exons (n=4 each group). **B.** Boxplot showing differential expression of an Fn1 transcript containing EDA/EDB exons in ciDKD podocytes. **C.** Exon/intron structure of Fn1 transcript differentially expressed in ciDKD vs wild-type podocytes. Expression of an EDA and EDB exon-containing transcript (not normally expressed in mature podocytes and associated with fibrosis) is increased in ciDKD podocytes. **D.** qPCR shows decreased expression of Hcfc1r1 mRNA in ciDKD podocytes. *t*-test, ****p<0.00001, n=4 independent experiments. **E.** qPCR using primers specific to the Fn1 EDB exon shows an increase in the expression of fibrosis associated EDB exon-containing transcripts in ciDKD podocytes. *t*-test, **p<0.001, n=4 independent experiments.

### Multiple spliceosomal-protein post-translational modifications (PTM) occur in response to insulin and IGF1 stimulation

As our data demonstrated the spliceosome complex was disrupted in ciDKD podocytes, we used phosphoproteomics to establish if there were any insulin or IGF1 stimulated post-translational modifications (PTM) in our cells. Wild-type (WT) and ciDKD cells were studied 3 days after Cre application (Figure Supp 2B). Cells were fasted of insulin and IGF1 for 4 hours and then stimulated with either 10 nM insulin or 10 ng/ml IGF1 for 10 minutes before performing TMT based phosphoproteomics. PTM phosphorylation events were filtered for spliceosomal and associated proteins by cross-referencing with the 199 proteins found in R-MMU-72203 reactome pathway data base (mouse processing of capped intron-containing pre-mRNA) (Supplementary Table 5). 94 of these proteins appeared in our data set. We then determined those PTM that significantly changed between the receptor replete (WT) and deplete cells (ciDKD) at a p-value <0.005 in response to insulin or IGF1 stimulation. As there was little difference when stimulating cells with either insulin or IGF1 we combined the data. 70 statistically significant phosphorylation events were detected in 27 spliceosomal proteins (Figure 7A). A recent study ^37^ by Turewicz *et al* has comprehensively examined the dynamic phosphorylation status of human myotubes exposed to 100 nM of insulin from 1 and 60 minutes. It detected 49 insulin-stimulated spliceosomal phosphorylation events and demonstrated that 100 nM insulin rapidly altered the mRNA splicing pattern of these cells. We detected the same phosphorylation events in four of the proteins in our data set. Splicing factor 3A subunit 1 serine 329 (SF3A1[S329]) was present in both data sets; Serine/arginine-rich splicing factor 2 serine 189 and 191 (SRSF2[S189 S191] was altered in our data and SRSF2[S191] in Turewicz’s; Serine/arginine-rich splicing factor 1 serine 199 (SRSF1[S199]) appeared in our data and SRSF1[S199 S201 S205] in Turewicz’s. Finally, we detected a phosphorylation event(s) in heterogeneous nuclear ribonucleoprotein D (HNRNPD) located between amino acids 68-85 (Figure 7A). In Turewicz’s data HNRNPD was phosphorylated at S80 S83 in response to high dose insulin. Spliceosomal proteins throughout the complex were modified in response to insulin and IGF1 (Figure 7B). Some proteins exhibited multiple phosphorylation effects, including serine/arginine repetitive matrix protein 2 (SRRM2) which was modified extensively with 29 different PTM detected (Figure 7C). Considering this we went back to our phosphoproteomic data set and investigated if any kinases known to phosphorylate splicing factors were modulated by insulin/IGF1 and found that cyclin-dependent kinase 11 (Cdk11[S578]), serine/arginine-rich splicing factor protein kinase 1 Srpk1 [S311] were. Additionally, we discovered that protein-kinase C and casein kinase substrate in neurones 3 pacsin3 (S383) phosphorylation was significantly reduced in ciDKD cells (Figure Supp 5). Pacsin3 is a casein kinase 2 (CK2) substrate.

**Figure 7.**
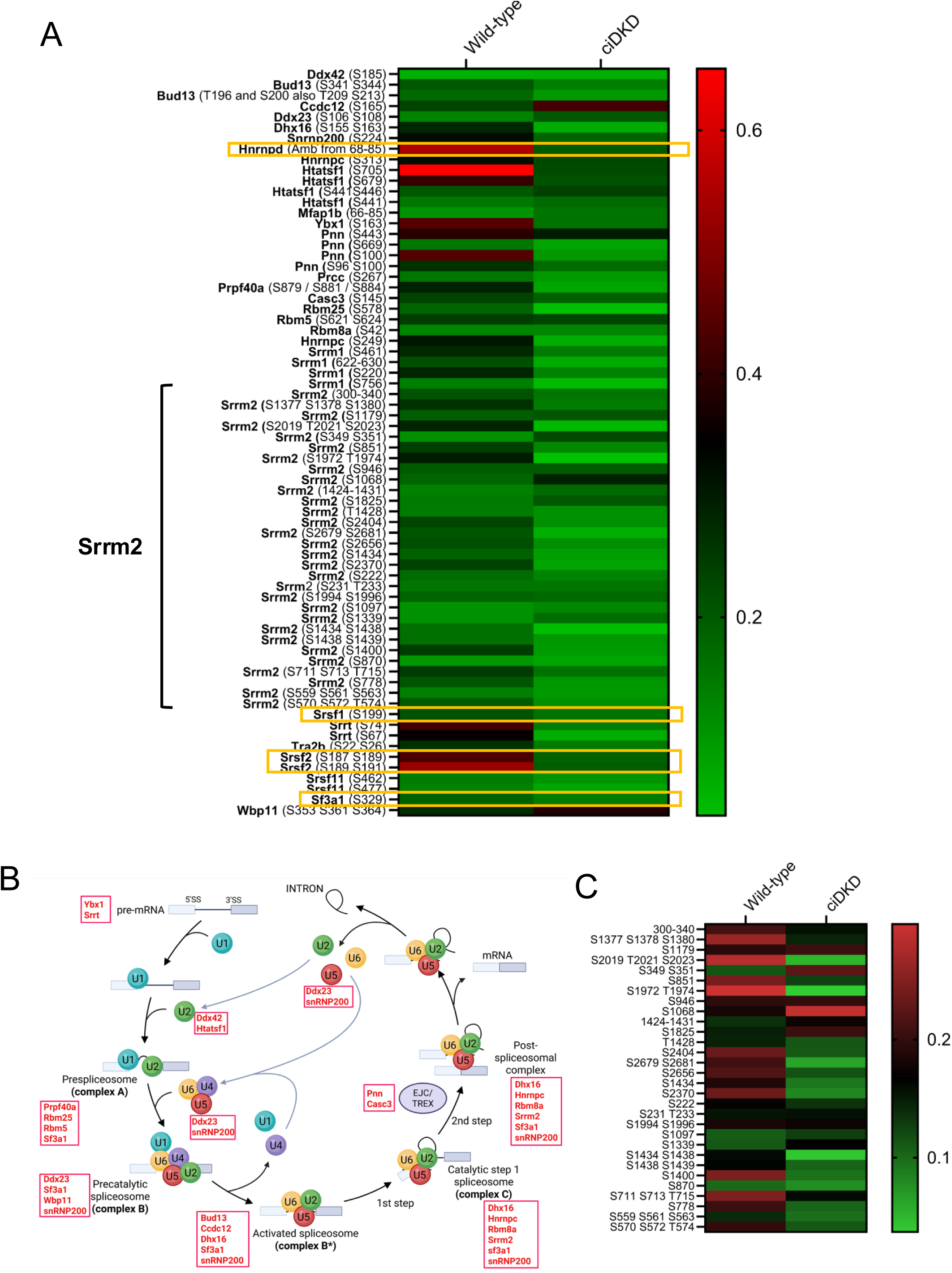
Multiple spliceosomal-protein post-translational modifications (PTM) occur in response to insulin and IGF1 stimulation. **A.** Heat map showing differential phosphorylation at the indicated phosphosites of spliceosome-related proteins in ciDKD podocytes and control cells stimulated with 10 nM insulin or 10 ng/ml IGF1 for 10 minutes (decreased phosphorylation in green, increased phosphorylation in red). Those phosphorylation events that also occurred in Turewicz study [37] highlighted in yellow boxes. **B.** Differentially phosphorylated proteins identified in A occur throughout the spliceosome cycle (red boxes). **C.** Heat map showing differential phosphorylation at the indicated phosphosites of serine/arginine repetitive matrix 2 (Srrm2) in ciDKD podocytes and control cells stimulated with 10 nM insulin or 10 ng/ml IGF1 for 10 minutes (decreased phosphorylation in green, increased phosphorylation in red).

### Relative contributions of the IR and IGF1R to spliceosomal modulation

To investigate the relative contributions of the IR and IGF1R to spliceosome control, we also generated conditionally immortalised podocyte cell single-receptor knock-down lines lacking either the IR [ciIRKD] or the IGF1R [ciIGF1RKD] (Figure Supp 6A and Supp 6B) in isolation and performed total and phosphoproteomic analysis on these and compared to the ciDKD cells. For total proteome analysis Perseus software was used to compare the four podocyte cell lines (wild-type, ciIRKD; ciIGF1RKD and ciDKD). Principal component analysis determined that samples were clustered according to experimental group (Figure Supp 6C-E) and a heat map with hierarchical clustering of proteins was subsequently generated to compare the four experimental groups (Figure 8A). The spliceosome targets identified from proteomic analysis of WT vs ciDKD podocytes, and which were validated by western blotting (Figure 3), were mapped on to this new heat map (Figure 8A). This revealed that while Eif4a3 and Ptbp2 downregulation were restricted to ciDKD and ciIGF1RKD podocytes, reduced Sf3b4 expression occurred in all three knockout cell lines. When compared to wild-type cells, Sf3b4 expression was lowest in ciDKD podocytes (fold change −0.56) while in ciIGF1RKD and ciIRKD the fold change was −0.28 and −0.34 respectively. Functional enrichment analysis comparing the four cell lines was performed and identified enrichment of the Gene Ontology term; spliceosomeal tri-snRNP complex assembly exclusively in the ciDKO podocytes (Figure 8B).

**Figure 8.**
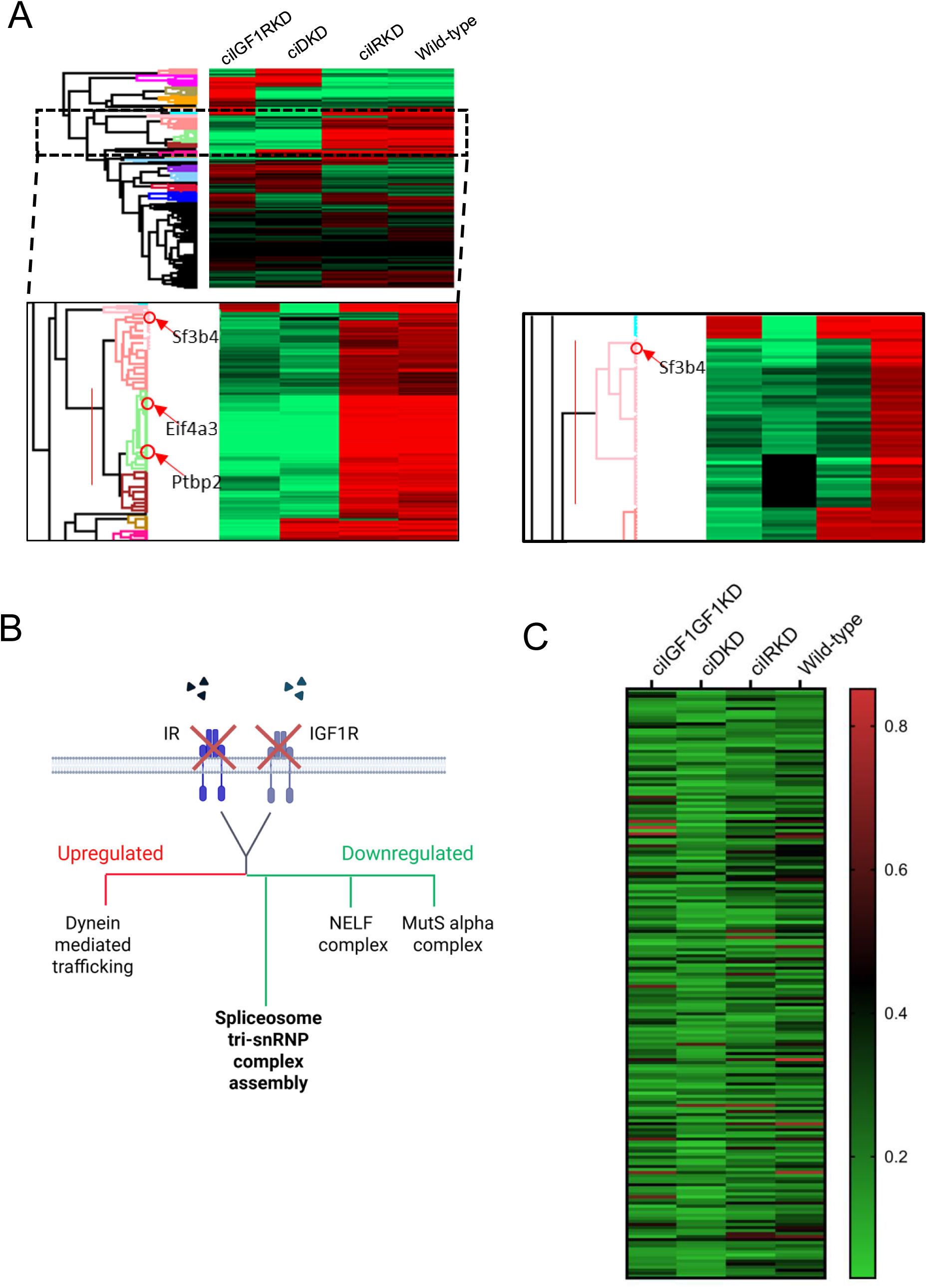
Relative contributions of the IR and IGF1R to spliceosomal modulation. **A.** Heatmap of hierarchical clustering of ciDKD vs ciIGF1RKD vs ciIRKD vs wild-type podocytes (n=9) proteomes (high relative expression in red, low relative expression in green, equal expression in black). Top of the heat map contains most significantly enriched groups, with enrichment intensity decreasing further down. The spliceosomal targets Sf3b4, Eif4a3, and Ptbp2 were mapped onto the proteomic heatmap comparing wild-type podocytes to cells with knockdown of the IR and/or the IGF1R. Sf3b4 expression is reduced in all three knockdown cell lines. **B.** Schematic showing signaling pathways uniquely enriched in ciDKD podocytes including the downregulation of spliceosomal tri-snRNP complex assembly. **C.** Heat map showing differential phosphorylation events in ciIGF1RKD; ciDKD; ciIRKD and wild-type podocytes (decreased phosphorylation in green, increased phosphorylation in red).

To assess the relative contribution of each receptor to spliceosomal PTM modulation we re-analysed our phosphoproteomic data sets comparing ciDKD cells with single receptor knockout lines stimulated with either 10 nM insulin or 10 ng/ml IGF1 for 10 minutes. We found that 49/70 PTM had signal in all cell lines studied (Figure 8C). There was no consistent patten of a single receptor being more dominant in this analysis. Some modifications were predominant through the IR [Ddx42(S185), Srsf11(S462), Srrm2(S1400)], some through the IGF1R [Srrm2 (S222, S349, S351, S570 S572 T574, S778, S870 S1825, T1428 S1434, S1438, S1439, S2370, S2679, S2681), Casc3 (S145), Htatsf1 (S441)] but most were partially abrogated through both. Collectively, this data suggests a complex network of insulin and IGF signalling via both receptors modulating the spliceosome complex.

## DISCUSSION

We have previously shown that signalling to the podocyte via the IR is critical for kidney function ^17^ while the IGF1R has a role in maintaining podocyte mitochondrial function ^21^. The current study used *in vivo* and *in vitro* models lacking both receptors simultaneously to further elucidate important common signalling pathways regulated by the insulin/IGF1 axis in the podocyte.

We initially used a transgenic mouse model (podDKD) to determine the physiological consequences of dual receptor loss *in vivo*. These mice developed an obvious renal phenotype: significant albuminuria, glomerulosclerosis, and ultrastructural filtration barrier damage was apparent by 6 months and was severe enough to cause death in several mice. However, body weight and blood glucose in podDKD mice were comparable to controls, in contrast with several other mouse models where alongside tissue-specific manifestations, double receptor knockout also gives rise to systemic effects. For example, IR/IGF1R knockout in kidney proximal tubule cells ^10^ and muscle ^11^ both result in decreased body size while other models have impairments in systemic metabolism: mice with adipose tissue-specific IR/IGF1R knockout develop diabetes ^9^ while dual receptor knockout in hippocampus and central amygdala results in a decrease in glutamate receptors and reduced glucose tolerance ^12^. Our models showed a variable phenotype which we think was predominantly due to the variable levels of receptor knockdown. We suspect if the IR and IGF1R knockdown had been >80% for both in the mice that a significant embryological / neonatal renal phenotype would have been observed as the podocin cre driver becomes active form approximately day 12 of development. For this reason, we subsequently generated podocyte cell lines in which we could reproducibly knock down the receptors by more than 80% to explore the underlying mechanism of disease.

We performed unbiased proteomic analysis to identify processes responsible for the detrimental phenotype observed in our models. This revealed major changes in the spliceosome complex. The spliceosome is a large ribonucleoprotein complex whose critical function in all eukaryotic cells is to catalyse the removal of introns from precursor messenger RNA molecules and join the flanking exons to generate mature mRNA transcripts that direct protein synthesis during translation ^38^. Spliceosome function is also a driver of significant proteomic diversity via alternative splicing, whereby multiple mRNA transcripts containing different exon combinations are produced from a single gene ^38^. Mutations in core spliceosome components, spliceosome assembly factors or regulatory proteins can result in defects in pre-RNA processing, increased nonsense-mediated decay of affected transcripts, abnormal or truncated protein products and changes to alternative splicing. Such erroneous splicing can substantially alter the transcriptome in specific cell types. This has been shown in immune cell dysfunction ^39^, cancer ^39^ and neurodegeneration ^40,41^. Somatic mutations in genes encoding spliceosome components can result in diseases, known as spliceosomopathies, (including retinitis pigmentosa, myelodysplastic syndromes and craniofacial disorders ^42^), with tissue-specific manifestations, suggesting that although spliceosome factors are ubiquitously expressed, they likely have cell type specific functions. Interestingly, one of the splice factors frequently involved in these spliceosomopathies - SF3B4 - was down regulated in the double knock-down podocytes (Figure 3E) and in our murine model (Figure 3F). We used the spliceosome inhibitor pladienolide B ^43^, which targets the SF3B complex, to investigate the effect of reduced spliceosome activity in wild-type podocytes. Developed as an anti-neoplastic agent, pladienolide B is effective at reducing cell viability, particularly in cancer cell lines ^43–46^, presumably due to an increased demand for translation and protein production in such actively dividing cells. In contrast, a comparatively high dose of pladienolide B is not sufficient to induce cytotoxicity in HEK 293T cells, suggesting that not all cell types are equally sensitive to spliceosome inhibition ^47^. While we were able to reproduce the low dose cytotoxic effect of this spliceosome inhibitor in HeLa cells reported previously ^43^, the response of the glomerular cell types we assayed differed. No cytotoxic effect was apparent in glomerular endothelial cells while in contrast, we found podocytes to be sensitive to pladienolide B, resulting in significant cell death and suggestive of the importance of adequate spliceosome function in this cell type. We therefore performed long-read RNA sequencing to explore the global effect of podocyte IR/IGF1R loss on the transcriptome of ciDKD cells.

Long-read RNA Seq analysis showed that podocyte IR/IGF1R depletion had a marked impact on multiple alternative splicing events, notably an increase in the fraction of transcripts with intron retention/premature termination codons. Interestingly, enrichment analysis showed that many transcripts associated with spliceosome function and RNA transport had increased levels of intron retention, cassette exon splicing and alternative 5’ and 3’ splicing. Although it has recently been recognised that intron retention can serve as a mechanism for regulating gene expression ^48^ including the regulation of RNA processing genes ^49^, our results could also be explained by the apparent loss of the spliceosome in ciDKD podocytes. Indeed, intron retention secondary to mutations in spliceosome components is a hallmark of a number of cancers ^50^ and neurodegenerative conditions ^40^. Moreover, it is likely that our RNA seq analysis underestimates the incidence of transcripts with premature termination codons as many will be targeted for nonsense-mediated decay ^51^ and would thus be unmeasurable in our samples.

Significant differential transcript expression was apparent in ciDKD podocytes with transcripts encoding the extracellular matrix protein fibronectin among the top hits. We detected increased expression of a Fn1 splice variant which included the EDA and EDB exons. Although EDA/EDB-fibronectin is important during development, its expression is limited in adult tissue including kidney ^52^. However, EDA-fibronectin is upregulated in cultured podocytes exposed to TGFβ treatment ^53^ and a pathological increase in this isoform results in insulin resistance ^54^, together with fibrosis in multiple tissues including lung ^55^ and kidney ^56^. It is conceivable that the changes in podocyte splicing resulting from IR/IGF1R knockdown promote a fibrotic phenotype by altering the ratio of fibronectin variants expressed, consistent with the increased fibrosis observed in our podDKD mice. However, it is likely many detrimental biological processes are initiated through spliceosomal dysregulation.

Our phosphoproteomic data suggests there are multiple insulin and IGF1-mediated signalling events in a variety of spliceosomal proteins that cause post-translational modifications. This is supported by Turewicz’s recent work ^37^, which built on iPSC myoblast studies by Batista et al ^57^ showing that multiple spliceosomal PTM events occur in myotubes stimulated with 100 nM insulin at a variety of time points, ranging from 1 to 60 minutes. 100 nM insulin will signal through both the IR and IGF1R, as well as insulin/IGF1R hybrid receptors ^4^ which is relevant to our study. We discovered that some proteins appeared to have multiple insulin/IGF1 induced PTMs, most notably Srrm2 with 29 changes detected. Srrm2 is a large protein thought to act as a scaffold protein for nuclear speckles. Nuclear speckles function as reservoirs of splicing factors, coordinating their release into the nucleoplasm to regulate transcription ^58^. It has previously been shown that SRRM2 phosphorylation status is key for its spliceosomal function with nearly 80 phosphorylation sites identified ^59^. Due to these multiple phosphorylation events we looked for evidence that kinases known to phosphorylate spliceosome proteins were modified in response to insulin/IGF1 and found three kinases were differentially phosphorylated (CDK11[S578]), SRPK[S311], and PACSIN3[S383]). Interestingly PACSIN3 is a substrate for Casein Kinase 2 (CK2) which has recently been shown to be a key kinase in regulating SRRM2 ^59^.

We found a predictable downregulation of cell cycle and DNA replication pathways given the established mitogenic effect of IGF1 signalling ^60^. However, we also observed a decrease in DNA damage response pathways and histone methylation. As post-mitotic cells with a limited capacity for replenishment, maintaining genomic integrity is crucial for podocyte health ^61–63^. Furthermore, altered histone methylation, another feature of the podocyte IR/IGF1R knockdown proteome, has been shown to affect the function of the DNA repair system in podocytes ^64^. Recent studies suggest that podocyte DNA damage and changes in glomerular DNA methylation may be useful markers of chronic kidney disease and correlate with estimated glomerular filtration rate (eGFR) decline in patients with diabetic nephropathy ^65^. It is therefore plausible that deficiencies in DNA damage repair mechanisms also contribute to the detrimental phenotype and ultimate loss of podocytes resulting from IR/IGF1R knockdown.

Evidence from previous studies suggests that DNA damage is intrinsically linked with spliceosome function: DNA damage affects the expression, activity and localisation of splicing factors ^66^ while spliceosome dysfunction leads to DNA damage by promoting the aberrant formation of R loops during transcription ^67,68^. Thus, accurate spliceosome function is critical for maintaining genomic integrity ^67^. In addition to the decreased expression of DNA damage response proteins revealed by our proteomic data, we observed a striking reduction in the levels of proteins required for spliceosome assembly and regulation in ciDKD podocytes. We have previously used lentiviral mediated expression of Cre recombinase to induce knockout of other podocyte genes, notably glycogen synthase kinase 3 α and β in combination ^27^. While this resulted in a highly detrimental cellular phenotype, proteomic analysis of these cells showed no differences in the expression of spliceosome related proteins, so we are confident that the results from the current study are not due to generalised podocyte cell death/dysfunction.

Finally, we explored the relative contribution of the IR and IGF1R to spliceosomal dysfunction. From a signalling perspective, we discovered that there were some spliceosome protein PTMs predominantly induced through the insulin or IGF1 receptor. However, many PTMs required the loss of both receptors to be phosphorylated or dephosphorylated. This suggests a complex signalling cascade that has evolved to compensate for a single receptor loss. Our total proteomic analysis suggested perhaps a predominance of the IGF1 receptor for spliceosome function, but pathway enrichment analysis revealed tri-snRNP complex assembly, a key process in the spliceosome cycle, required for the formation of the pre-catalytic spliceosome complex B, was only down regulated when both receptors were depleted.

There are several limitations to our studies: Our mouse model showed variability which we think was predominantly driven by the variable level of receptor knockdown but may have also been the result of studying mice on a mixed genetic background (as we consider this more relevant to human disease). We could have studied a single background (or backgrounds) to explore this. Furthermore, we could have investigated the early effects of partial receptor knockdown in the embryonic / neonatal phase in the mice. We also considered an inducible cre system to study IR/IGF1R loss in fully developed adult podocytes but these systems result in much less gene excision, as shown by our reporter studies, and would likely have had a very mild or no phenotype.

For our *in vitro* work we only examined a single time point (10 minutes) and dose of insulin and IGF1 in our phosphoprotein omics studies. We think a more comprehensive dose-time/course of insulin/IGF1 phosphorylation changes may reveal more detail of the signalling pathways induced through the IR and IGF1R in isolation and together. The doses of insulin and IGF1 used are likely not specific for their cognate receptor and the kinetics of phosphorylation events probably spans for minutes to hours in this system as shown in the study of insulin in myotubes ^57^. It may also have been the case that some of the PTMs we discovered are the result of evolving damage to the spliceosome, which could have been induced or exacerbated by the putative DNA damage revealed by our proteomic analysis. Future work will include a comprehensive time and dose response analysis of WT and receptor deficient cells to more precisely delineate the important signalling pathways in this process. Additionally, this may identify targets for genetic manipulation to see if we can rescue the phenotype.

In summary, we have shown that the IR/IGF1R axis is critical for podocyte health. Mechanistically, this axis controls gene transcription by maintaining spliceosome function. This suggests hormonal signals are important for the genetic control of these terminally differentiated cells.

## Acknowledgements

We thank the Proteomic Facility and Histology Services Unit at the University of Bristol for technical support. This study was funded by the Medical Research Council (Senior Clinical Fellowship) to RJMC-(MR/K010492/1), Kidney Research UK (RP_011_20230628), and funding from the Innovative Medicines Initiative 2 Joint Undertaking (JU) under grant agreement No 115974. The JU receives support from the European Union’s Horizon 2020 research and innovation programme and EFPIA and JDRF. Any dissemination of results reflects only the author’s view; the JU is not responsible for any use that may be made of the information it contains.

## Author contributions

The study was conceived by R.J.M.C., G.I.W. and J.A.H. Latterly academic input was supplied by M.H. and P.T.B. Experiments were performed by J.A.H., L.D., J.T.C. and F.B. Conditionally immortalised cell lines were made by L.N. PodCre mice were provided by P.T.B. IGF1R^fl/fl^ mice were provided by M.H. Long-read RNA sequencing was performed and processed by A.J. and S.B., F.B. and M.I. bioinformatically analysed and interpreted the data. The manuscript was written by J.A.H and R.J.M.C. and all authors reviewed, commented and approved the paper.

---

**Figure S1.**
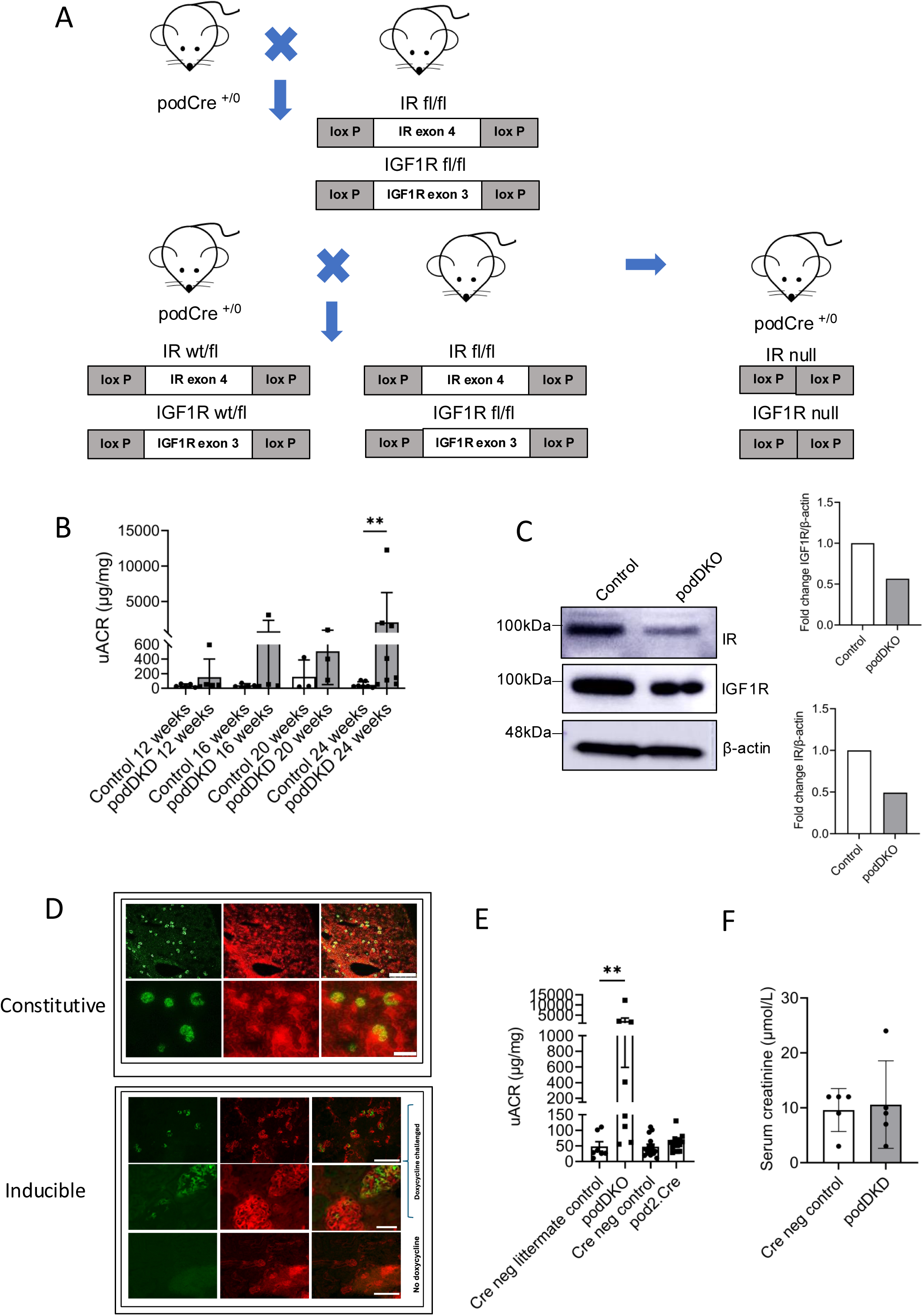
Generation of podDKO mice. Related to Figure 1. **A.** Breeding scheme used to generate podDKD mice. **B.** Progression of albuminuria in podDKD mice. Showing that uACR becomes significantly increased by 24 weeks. Unpaired t test, **p<0.005. **C.** Western blot of primary podocytes derived from podDKD and Cre negative littermate control mice **D.** Top panel – constitutive podocin cre mouse (het) crossed with mTR/mGFP reporter. Cre expression causes GFP to be switched on (>80% excision). Bottom panel – Inducible Podocin RtTA –tet-o-cre after 14 days doxycycline (approximately 50% excision). Scale bar either 250 micrometres (high magnification) or 50 micrometres (low -magnification) **E.** No difference in uACR between Cre negative podDKD littermate controls, Cre negative and pod2.Cre expressing mice at 6 months. uACR is significantly increased in podDKD mice. *t-*test, **p<0.01, n=5-8 mice per group. **F.** Graph showing serum creatinine levels in subset of control and podDKD mice age (n=5 each group).

**Figure S2.**
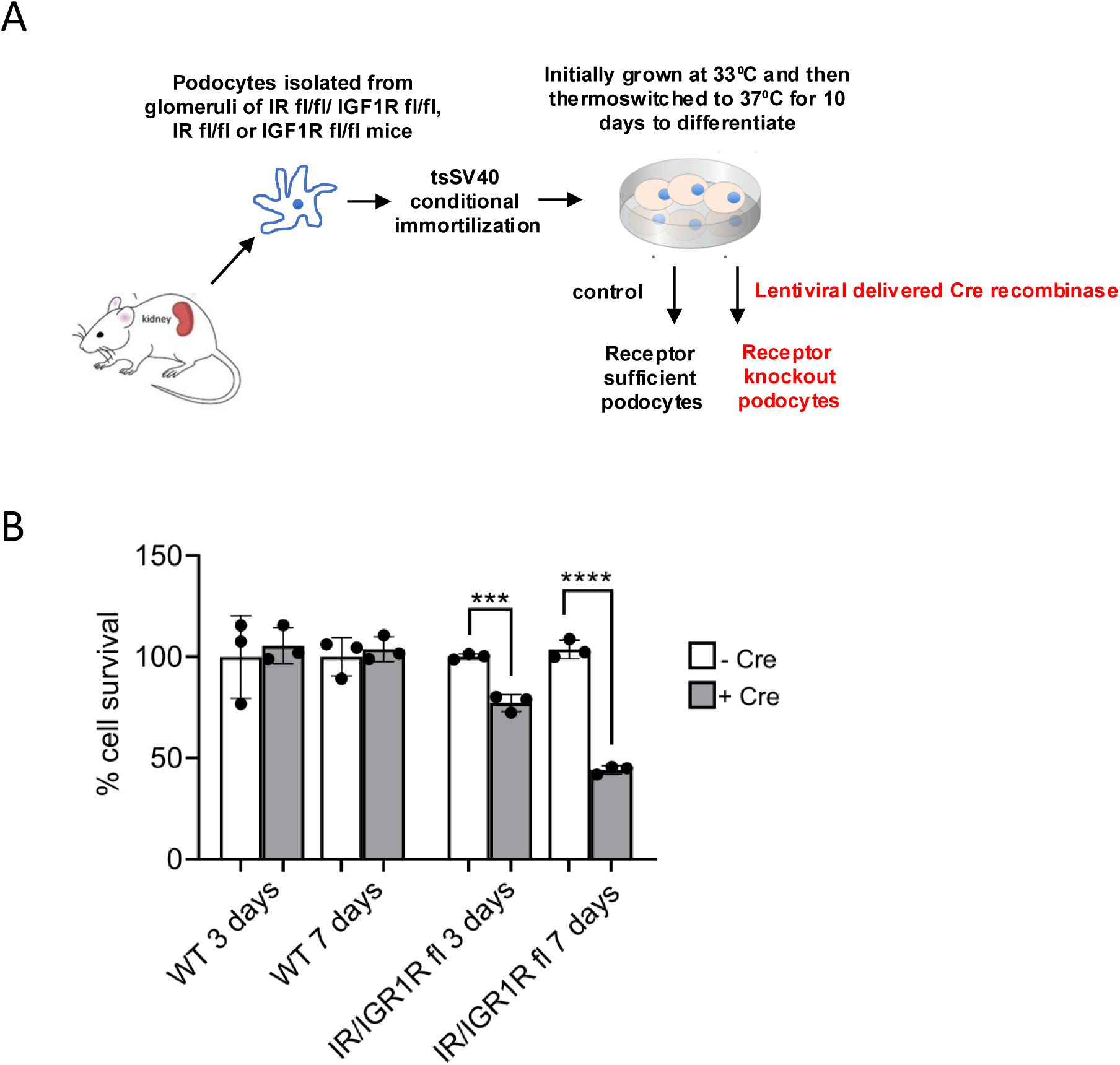
Related to Figure 2. **A.** Development of ciDKD, ciIRKD and ciIGF1RKD podocytes. Primary culture podocytes (homozygous floxed) isolated from transgenic mouse. Conditional immortalisation with temperature sensitive SV40 construct and then excision of IR and/or IGF1R using lentiviral delivered Cre recombinase. **B.** Survival rate of Wild-type and IR/IGF1R floxed podocytes 3 days and 7 days after Cre recombinase induced gene knockdown.t test, ***p<0.001, ****p<0.0001. (n=3 each group)

**Figure S3.**
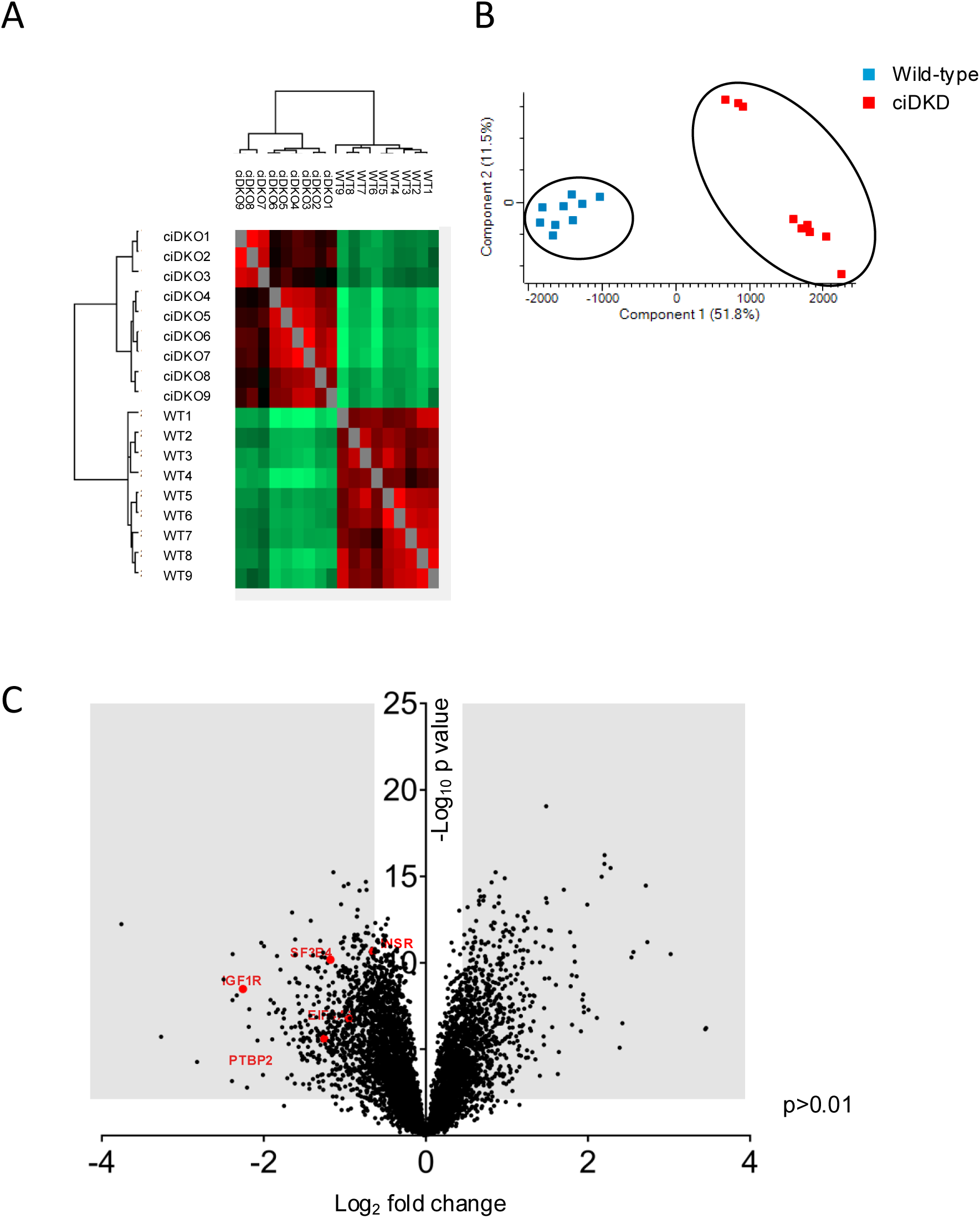
Proteomic analysis of ciDKD podocytes. Related to Figure 3. **A.** Heat map of sample clustering **B.** Plot of principal component analysis **C.** Volcano plot showing changes in the proteome of ciDKD podocytes relative to wild-type control cells; log2 fold change (FC) vs -log10 p-value of the scaled abundances. Shaded areas show proteins with FC < and > 2.

**Figure S4.**
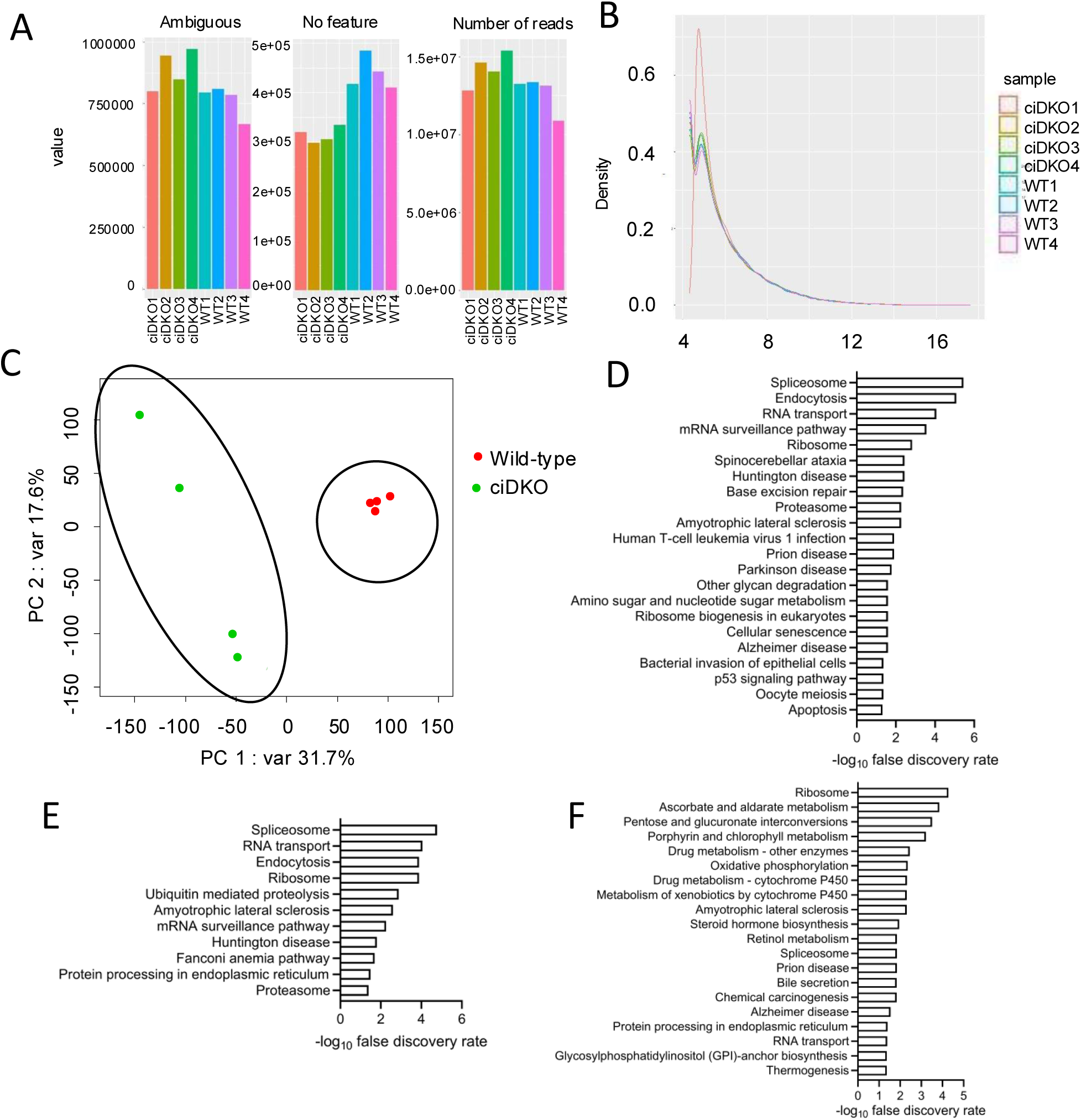
Long read RNA sequencing analysis of ciDKD podocytes. Related to Figures 5 and 6. **A.** Plot indicating the number of reads sequenced. **B.** Plot of variance transformed data to normalise for library size. **C.** Plot of principal component analysis shows clustering of experimental groups. **D-F.** KEGG enriched terms in ciDKD podocytes genes with intron retention (**D**), cassette exon splicing (**E**) and alternative 5’ splicing (**F**).

**Figure S5.**
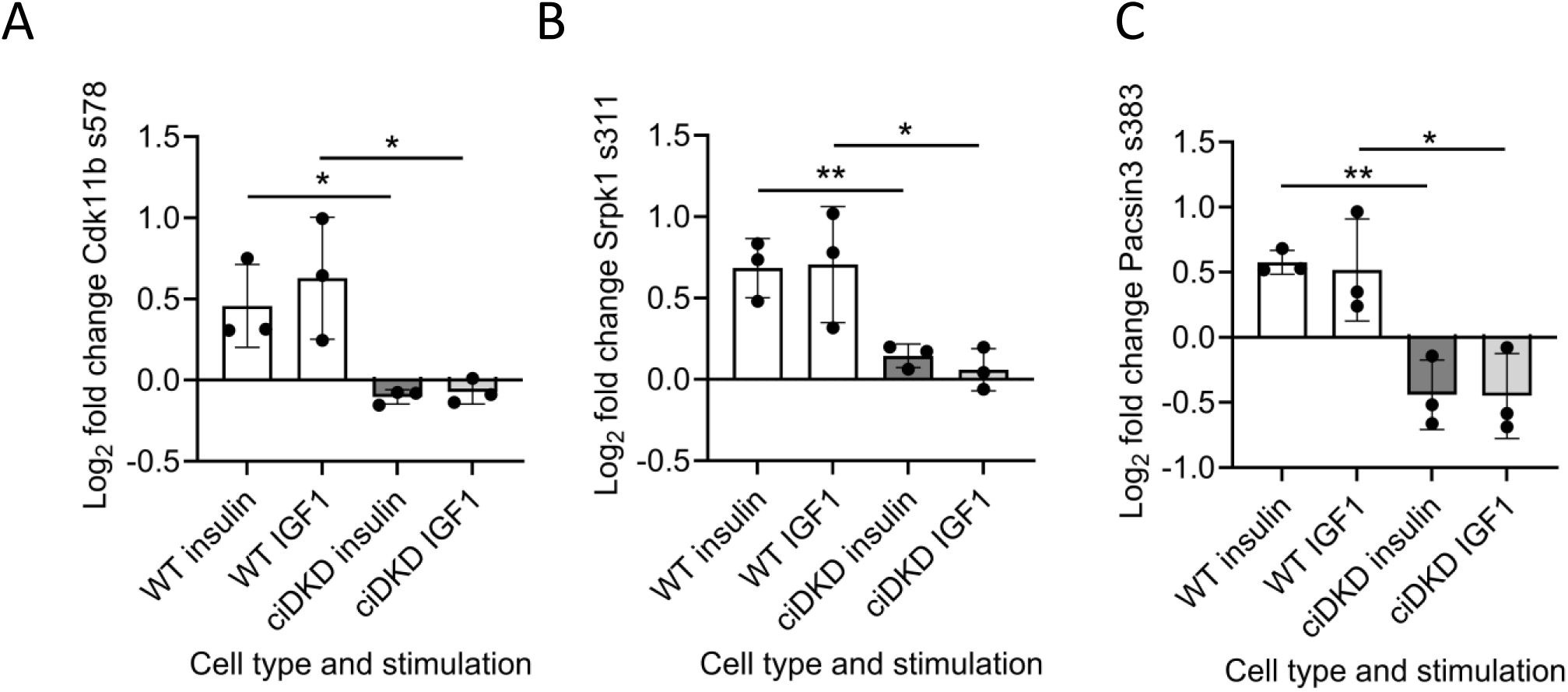
Spliceosomal-protein related kinase post-translational modifications (PTM) occuring in response to insulin and IGF1 stimulation. Related to Figure 7. Graphs showing the relative expression of Cdk11b s578 (**A**), Srpk1 s311 (**B**) and Pacsin3 s383 (**C**) in wild-type and ciDKD podocytes stimulated with 10 nM insulin or 10 ng/ml IGF1. Unpaired t test *p<0.05. **p<0.005, n=3.

**Figure S6.**
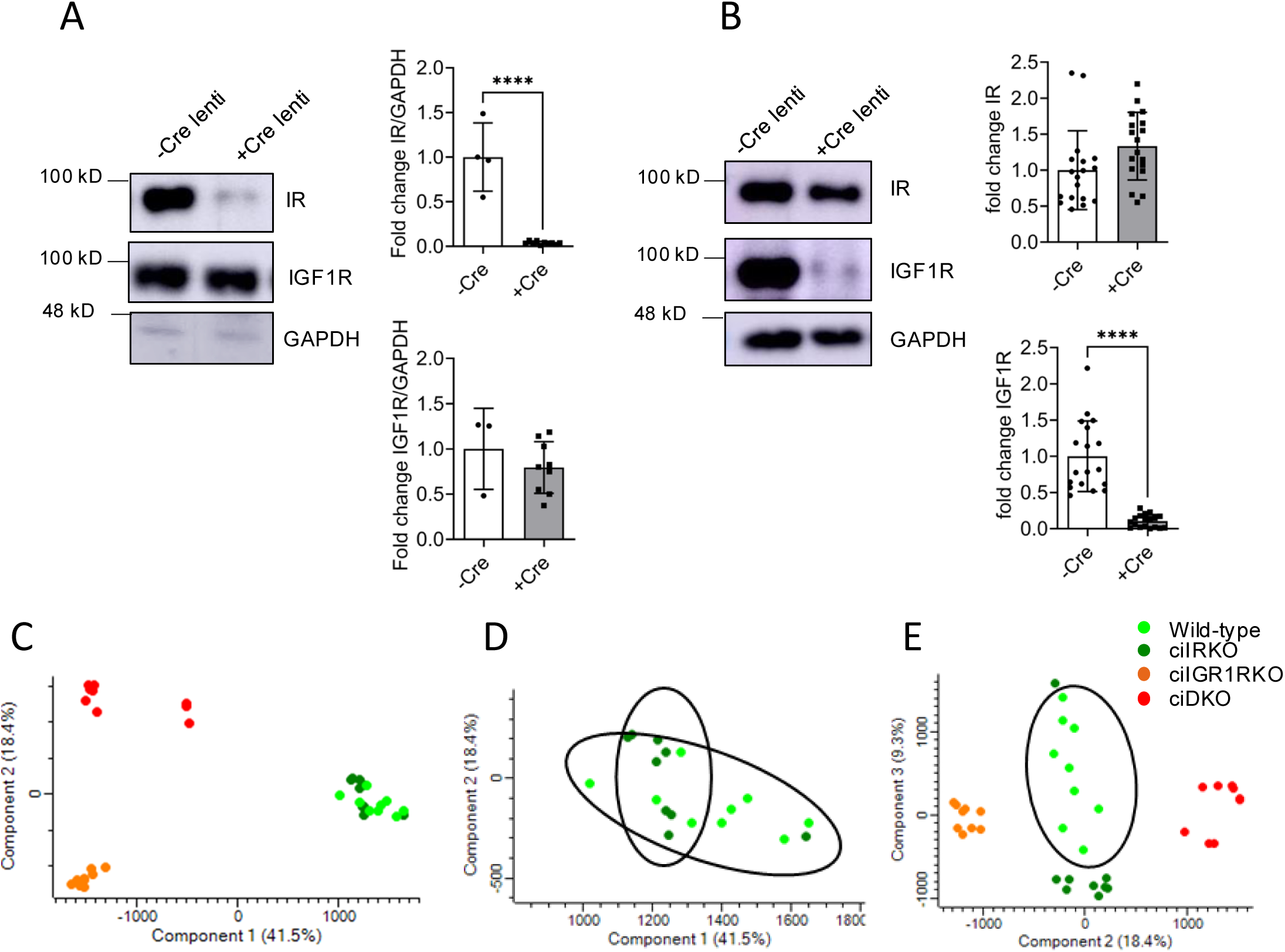
Proteomic analysis of ciIRKD, ciIGF1RKD and ciDKD podocytes. Related to Figure 8. **A.** Western blot shows >95% reduction of IR expression in ciIRKO podocytes, t test. ****p<0.0001. **B.** Western blot shows >95% reduction of IGF1R expression in ciIGF1RKD podocytes, t test. ****p<0.0001. **C-E**. Principal Component Analysis. ciDKD and ciIGF1RKD groups are significantly different to each other and the ciIRKD and wild-type groups **(C).** IRKD and WT groups show a moderate degree of overlap but do have a clear degree of difference**.(D).** 2D PCA analysis using third most significant component (Component 3), which demonstrates that along components 2 and 3, all 4 groups distinctly cluster separate to one another **(E)**.

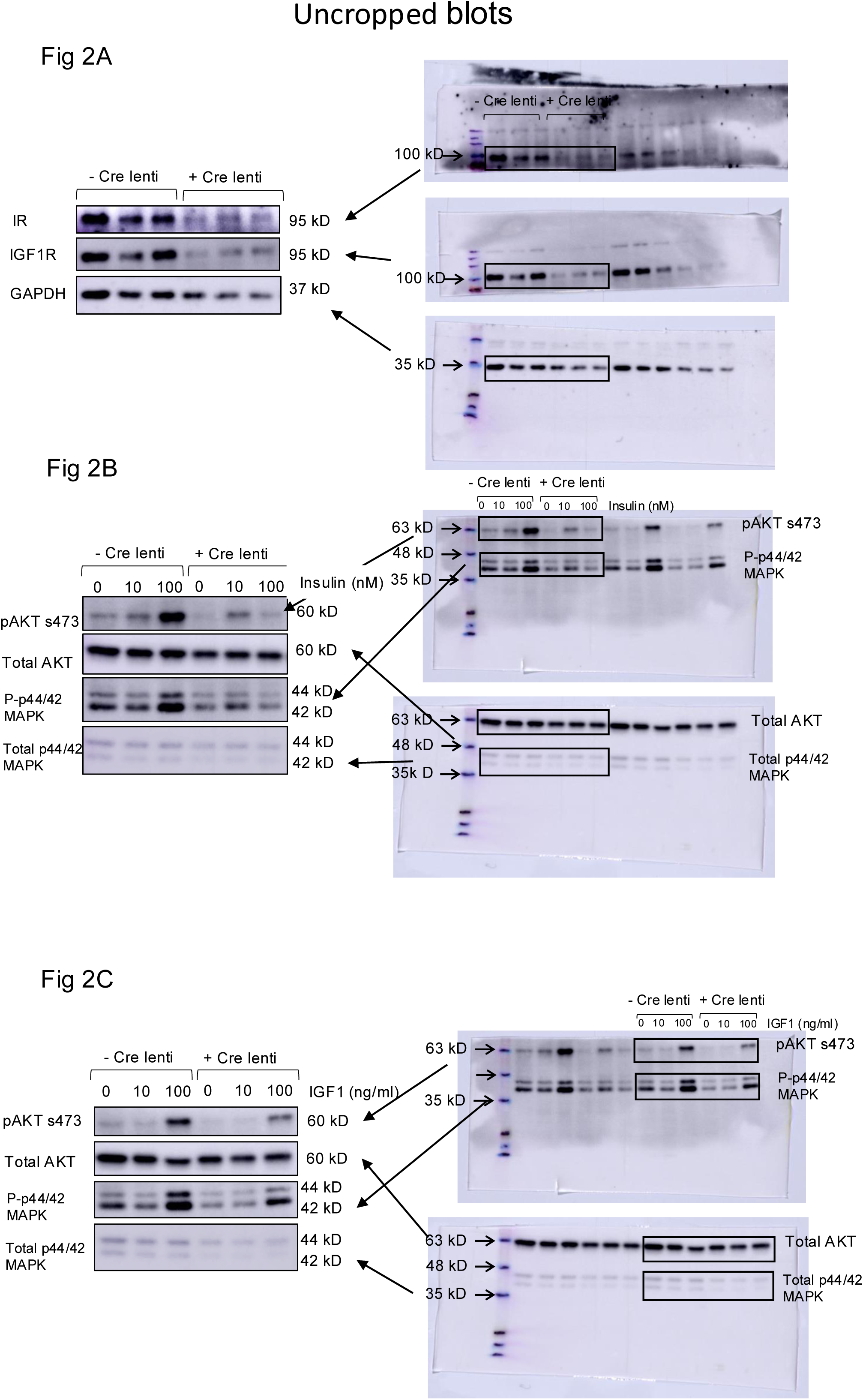

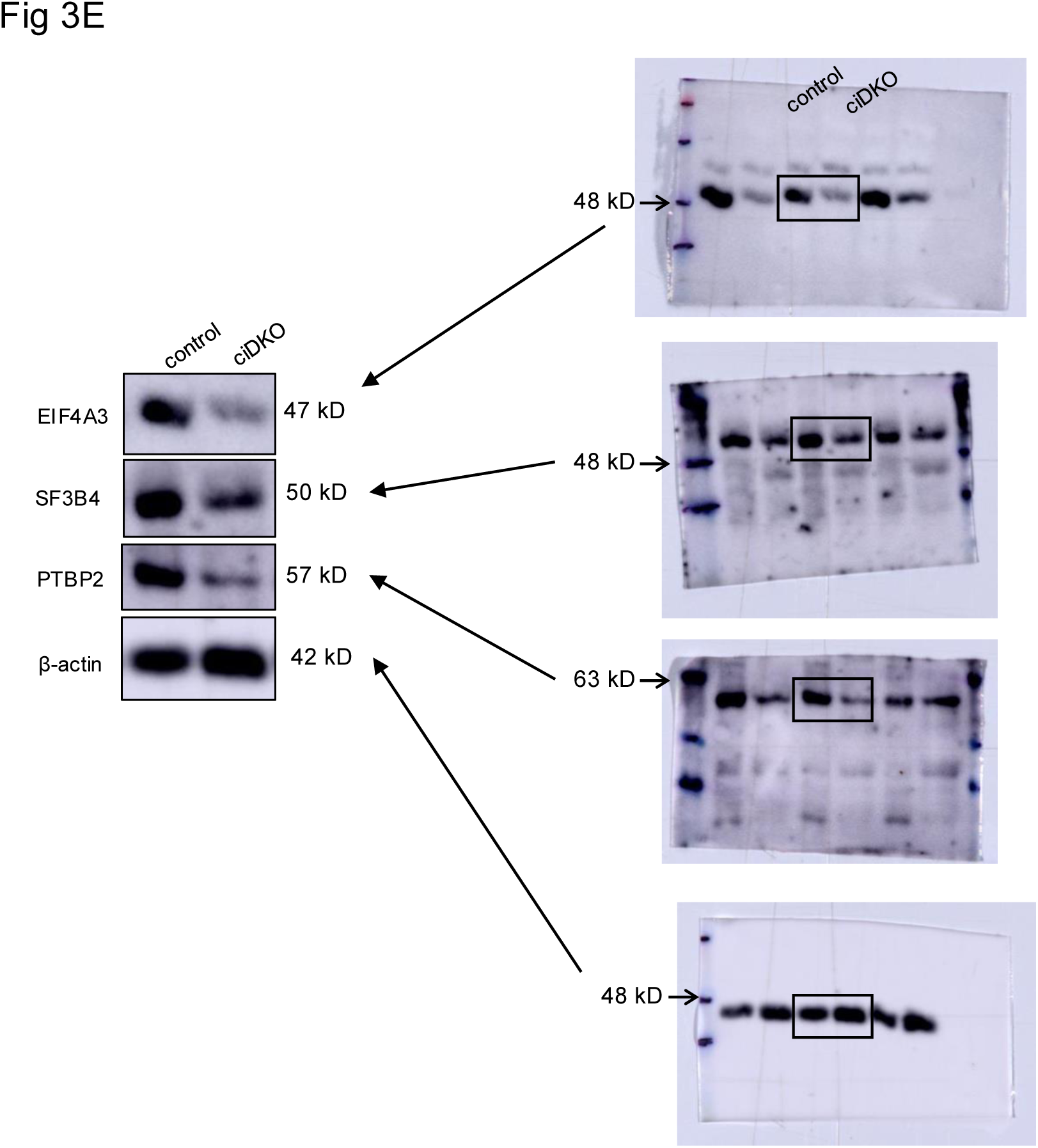

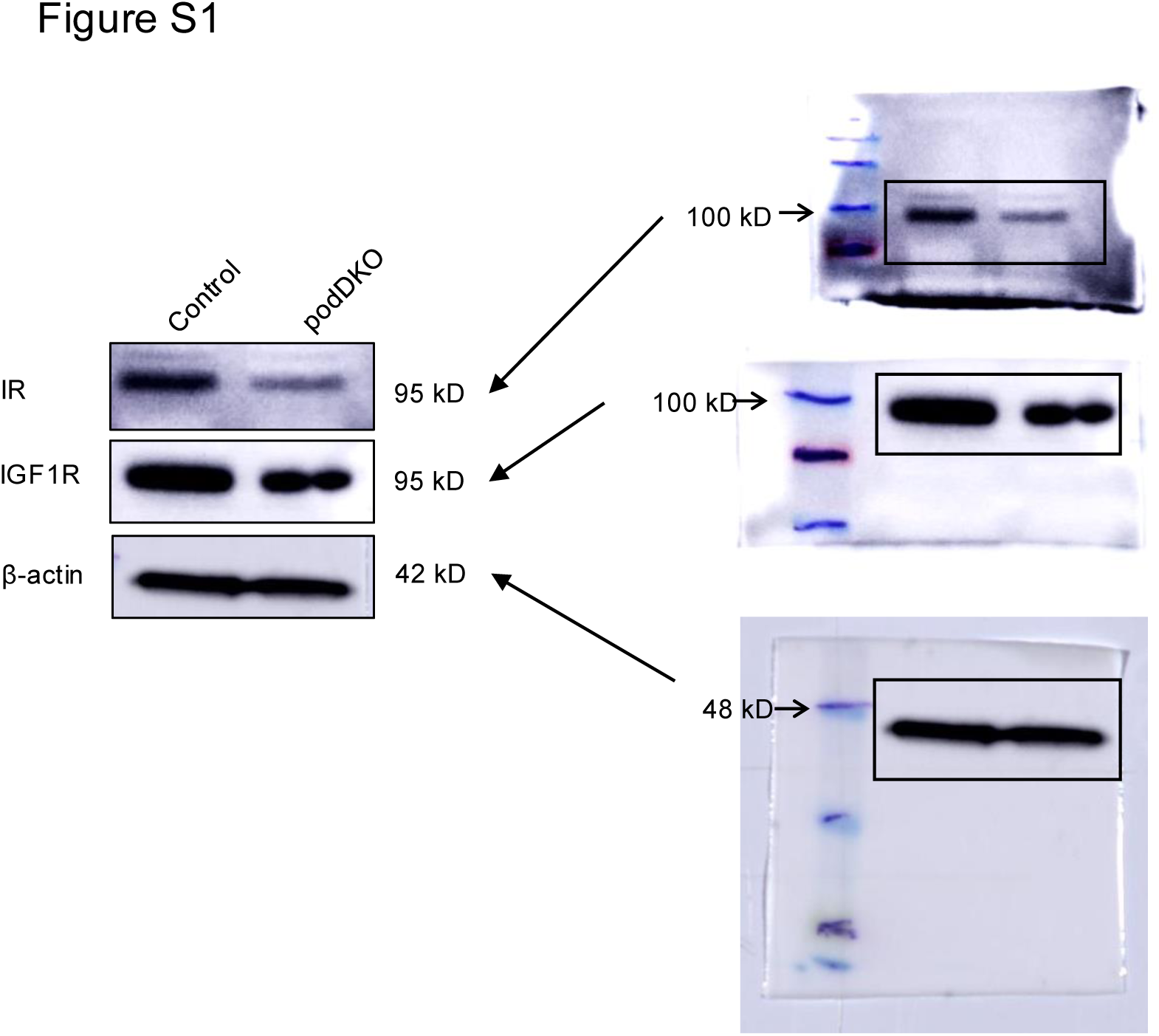

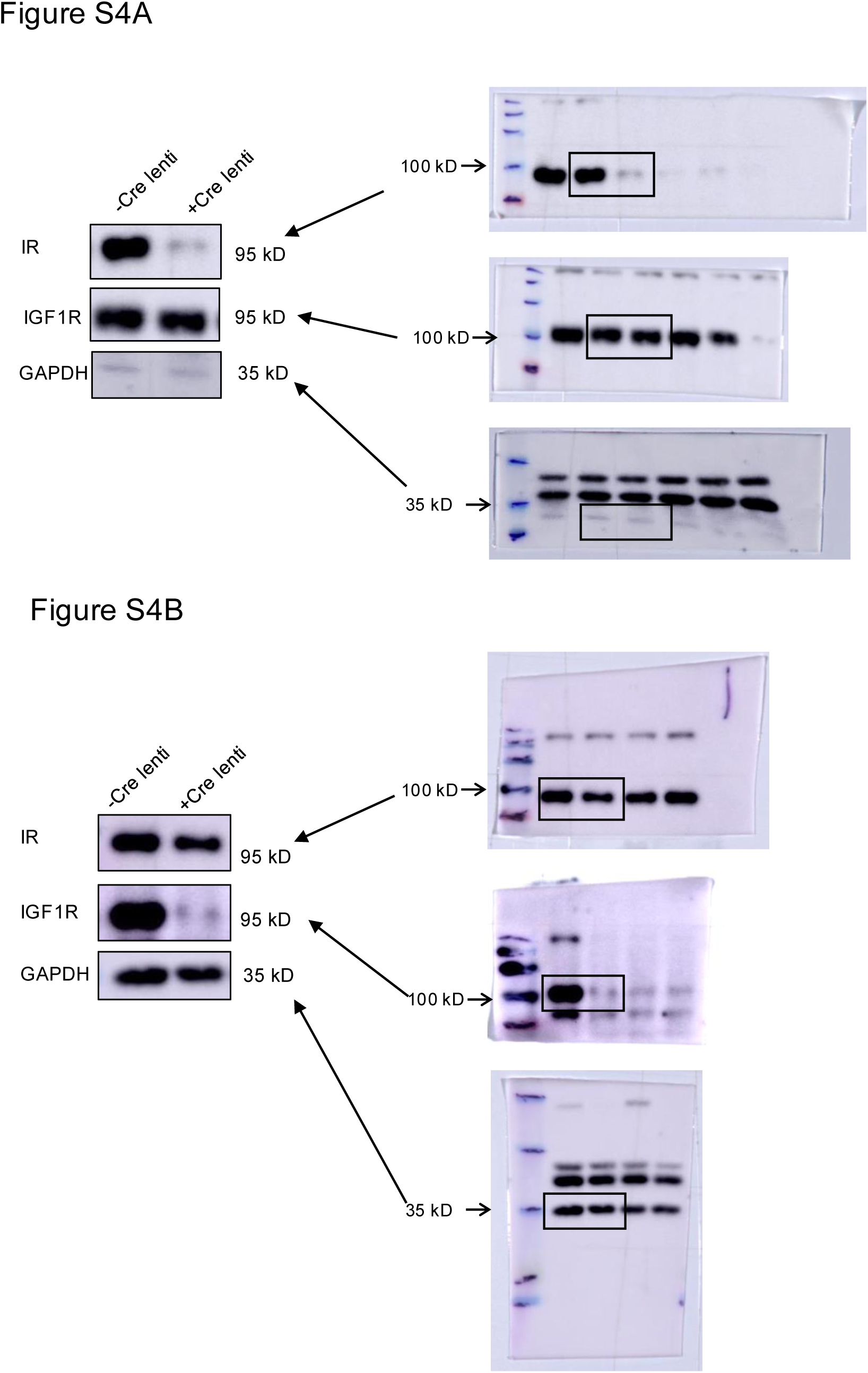

