## Supplementary material for "The insulin / IGF axis is critically important for controlling gene transcription in the podocyte": supp table 5 (from another version of this preprint: version 3, 2026-02-18)

| MoleculeType | Identifier | MoleculeName | |
|---|---|---|---|
| Chemical Cor | 30616 | ATP(4-) [ChEBI:30616] | |
| Chemical Cor | 456216 | ADP(3-) [ChEBI:456216] | |
| Chemical Cor | 43474 | hydrogenphosphate [ChEBI:43474] | |
| Chemical Cor | 18420 | magnesium(2+) [ChEBI:18420] | |
| Proteins | Q9CQ49 | UniProt:Q9CQ49 | |
| Proteins | Q3UYV9 | UniProt:Q3UYV9 | |
| Proteins | Q8R0A0 | UniProt:Q8R0A0 | |
| Proteins | Q3THK3 | UniProt:Q3THK3 | |
| Proteins | Q80UW8 | UniProt:Q80UW8 Polr2e | |
| Proteins | Q9D7M8 | UniProt:Q9D7M8 Polr2d | |
| Proteins | P61219 | UniProt:P61219 Polr2f | |
| Proteins | P62488 | UniProt:P62488 Polr2g | |
| Proteins | P08775 | UniProt:P08775 Polr2a | |
| Proteins | P97760 | UniProt:P97760 Polr2c | |
| Proteins | P62876 | UniProt:P62876 | |
| Proteins | O08740 | UniProt:O08740 Polr2j | |
| Proteins | Q923G2 | UniProt:Q923G2 Polr2h | |
| Proteins | P60898 | UniProt:P60898 Polr2i | |
| Proteins | Q63871 | UniProt:Q63871 Polr2k | |
| Proteins | Q8CFI7 | UniProt:Q8CFI7 Polr2b | |
| Proteins | P17225 | UniProt:P17225 | |
| Proteins | Q6PDM2 | UniProt:Q6PDM2 | |
| Proteins | Q9R0U0 | UniProt:Q9R0U0 | |
| Proteins | P84104 | UniProt:P84104 | |
| Proteins | Q8BL97 | UniProt:Q8BL97 | |
| Proteins | O88569 | UniProt:O88569 | |
| Proteins | Q9Z2X1 | UniProt:Q9Z2X1 | |
| Proteins | Q8VHM5 | UniProt:Q8VHM5 | |
| Proteins | Q8BG05 | UniProt:Q8BG05 | |
| Proteins | P70333 | UniProt:P70333 | |
| Proteins | Q60668 | UniProt:Q60668 | |
| Proteins | P61979 | UniProt:P61979 | |
| Proteins | P49312 | UniProt:P49312 | |
| Proteins | O35737 | UniProt:O35737 | also Q8C2Q7 |
| Proteins | Q8R081 | UniProt:Q8R081 | |
| Proteins | P56959 | UniProt:P56959 | |
| Proteins | Q9WV02 | UniProt:Q9WV02 | |
| Proteins | P62960 | UniProt:P62960 | |
| Proteins | P60335 | UniProt:P60335 | |
| Proteins | Q9Z204 | UniProt:Q9Z204 | |
| Proteins | Q8VEK3 | UniProt:Q8VEK3 Hnrnpu | |

| | | |
|---|---|---|
| Proteins | P62996 | UniProt:P62996 |
| Proteins | Q61990 | UniProt:Q61990 |
| Proteins | O35326 | UniProt:O35326 |
| Proteins | Q9D0B0 | UniProt:Q9D0B0 |
| Proteins | E9Q6E5 | UniProt:E9Q6E5 |
| Proteins | Q62093 | UniProt:Q62093 |
| Proteins | A2RRY9 | UniProt:A2RRY9 |
| Proteins | Q99KG3 | UniProt:Q99KG3 |
| Proteins | Q80X98 | UniProt:Q80X98 |
| Proteins | Q9DC48 | UniProt:Q9DC48 |
| Proteins | Q8BHJ9 | UniProt:Q8BHJ9 |
| Proteins | O08583 | UniProt:O08583 |
| Proteins | Q99M28 | UniProt:Q99M28 |
| Proteins | Q3ULL6 | UniProt:Q3ULL6 |
| Proteins | Q91VC3 | UniProt:Q91VC3 |
| Proteins | Q9CWZ3 | UniProt:Q9CWZ3 |
| Proteins | P61327 | UniProt:P61327 |
| Proteins | Q8K3W3 | UniProt:Q8K3W3 |
| Proteins | Q52KI8 | UniProt:Q52KI8 |
| Proteins | P26369 | UniProt:P26369 |
| Proteins | Q9D883 | UniProt:Q9D883 |
| Proteins | Q8BGJ9 | UniProt:Q8BGJ9 |
| Proteins | Q9Z1N5 | UniProt:Q9Z1N5 |
| Proteins | Q9CZU3 | UniProt:Q9CZU3 Mtrex |
| Proteins | P62311 | UniProt:P62311 |
| Proteins | Q9QXA5 | UniProt:Q9QXA5 |
| Proteins | P62322 | UniProt:P62322 |
| Proteins | P62313 | UniProt:P62313 |
| Proteins | Q9CQQ8 | UniProt:Q9CQQ8 |
| Proteins | Q6ZWM4 | UniProt:Q6ZWM4 |
| Proteins | O35900 | UniProt:O35900 |
| Proteins | Q9D0T1 | UniProt:Q9D0T1 |
| Proteins | Q922U1 | UniProt:Q922U1 |
| Proteins | Q9DAW6 | UniProt:Q9DAW6 |
| Proteins | Q9D868 | UniProt:Q9D868 |
| Proteins | P62320 | UniProt:P62320 |
| Proteins | P62307 | UniProt:P62307 |
| Proteins | P62309 | UniProt:P62309 |
| Proteins | P27048 | UniProt:P27048 |
| Proteins | P62317 | UniProt:P62317 |
| Proteins | P62315 | UniProt:P62315 |
| Proteins | P62305 | UniProt:P62305 |

| | | |
|---|---|---|
| Proteins | Q8CCF0 | UniProt:Q8CCF0 |
| Proteins | Q91V81 | UniProt:Q91V81 |
| Proteins | Q9Z315 | UniProt:Q9Z315 |
| Proteins | Q8K194 | UniProt:Q8K194 |
| Proteins | Q91YR7 | UniProt:Q91YR7 |
| Proteins | P83877 | UniProt:P83877 |
| Proteins | O08810 | UniProt:O08810 |
| Proteins | Q6PE01 | UniProt:Q6PE01 |
| Proteins | Q6P4T2 | UniProt:Q6P4T2 |
| Proteins | D3Z0M9 | UniProt:D3Z0M9 |
| Proteins | Q99PV0 | UniProt:Q99PV0 |
| Proteins | Q3TIX9 | UniProt:Q3TIX9 |
| Proteins | Q9R1C7 | UniProt:Q9R1C7 |
| Proteins | O70133 | UniProt:O70133 |
| Proteins | Q8BTI8 | UniProt:Q8BTI8 |
| Proteins | Q99MR6 | UniProt:Q99MR6 |
| Proteins | Q61656 | UniProt:Q61656 |
| Proteins | Q3UKJ7 | UniProt:Q3UKJ7 |
| Proteins | Q8JZX4 | UniProt:Q8JZX4 |
| Proteins | Q8BGT7 | UniProt:Q8BGT7 |
| Proteins | P57784 | UniProt:P57784 |
| Proteins | O35286 | UniProt:O35286 |
| Proteins | Q9CQI7 | UniProt:Q9CQI7 |
| Proteins | Q3UEB3 | UniProt:Q3UEB3 |
| Proteins | Q8CGZ0 | UniProt:Q8CGZ0 |
| Proteins | P63163 | UniProt:P63163 |
| Proteins | Q569Z5 | UniProt:Q569Z5 |
| Proteins | Q99NB9 | UniProt:Q99NB9 |
| Proteins | Q923D4 | UniProt:Q923D4 |
| Proteins | Q3UJB0 | UniProt:Q3UJB0 |
| Proteins | Q810A7 | UniProt:Q810A7 |
| Proteins | Q921M3 | UniProt:Q921M3 |
| Proteins | Q8QZY9 | UniProt:Q8QZY9 |
| Proteins | Q8K4Z5 | UniProt:Q8K4Z5 |
| Proteins | Q62203 | UniProt:Q62203 |
| Proteins | Q9D554 | UniProt:Q9D554 |
| Proteins | Q6NV83 | UniProt:Q6NV83 |
| Proteins | P83870 | UniProt:P83870 |
| Proteins | Q62189 | UniProt:Q62189 |
| Proteins | Q62241 | UniProt:Q62241 |
| Proteins | Q62376 | UniProt:Q62376 |
| Proteins | Q61136 | UniProt:Q61136 |

| | | |
|---|---|---|
| Proteins | Q91YE7 | UniProt:Q91YE7 |
| Proteins | B2RY56 | UniProt:B2RY56 |
| Proteins | Q9Z1M8 | UniProt:Q9Z1M8 |
| Proteins | Q8VH51 | UniProt:Q8VH51 |
| Proteins | Q8CGF7 | UniProt:Q8CGF7 |
| Proteins | O55128 | UniProt:O55128 Sap18 |
| Proteins | Q9JIX8 | UniProt:Q9JIX8 |
| Proteins | O35691 | UniProt:O35691 |
| Proteins | C0HKD8 | UniProt:C0HKD8 |
| Proteins | C0HKD9 | UniProt:C0HKD9 |
| Proteins | Q9EPV8 | UniProt:Q9EPV8 |
| Proteins | Q923D5 | UniProt:Q923D5 |
| Proteins | Q8BIZ6 | UniProt:Q8BIZ6 |
| Proteins | Q8R149 | UniProt:Q8R149 |
| Proteins | Q8R0F5 | UniProt:Q8R0F5 |
| Proteins | Q61048 | UniProt:Q61048 |
| Proteins | Q4FK66 | UniProt:Q4FK66 |
| Proteins | Q9CPW7 | UniProt:Q9CPW7 |
| Proteins | Q91VJ5 | UniProt:Q91VJ5 |
| Proteins | Q9D6L8 | UniProt:Q9D6L8 |
| Proteins | Q9D0W5 | UniProt:Q9D0W5 |
| Proteins | Q9EQC8 | UniProt:Q9EQC8 |
| Proteins | Q8R1N0 | UniProt:Q8R1N0 |
| Proteins | Q8CFQ3 | UniProt:Q8CFQ3 |
| Proteins | Q8BHS3 | UniProt:Q8BHS3 |
| Proteins | Q6PGH1 | UniProt:Q6PGH1 |
| Proteins | Q69ZQ2 | UniProt:Q69ZQ2 |
| Proteins | P63154 | UniProt:P63154 |
| Proteins | Q9CSN1 | UniProt:Q9CSN1 Snw1 |
| Proteins | Q9DCD2 | UniProt:Q9DCD2 |
| Proteins | Q8R344 | UniProt:Q8R344 |
| Proteins | Q3TKY6 | UniProt:Q3TKY6 |
| Proteins | Q8R3P8 | UniProt:Q8R3P8 |
| Proteins | Q9D787 | UniProt:Q9D787 |
| Proteins | G3X8X0 | UniProt:G3X8X0 |
| Proteins | P63017 | UniProt:P63017 Hspa8 |
| Proteins | Q9CWL8 | UniProt:Q9CWL8 |
| Proteins | Q9JHS9 | UniProt:Q9JHS9 |
| Proteins | Q99KP6 | UniProt:Q99KP6 |
| Proteins | Q9D287 | UniProt:Q9D287 |
| Proteins | Q922V4 | UniProt:Q922V4 |
| Proteins | Q6A068 | UniProt:Q6A068 |

| | | |
|---|---|---|
| Proteins | Q56A08 | UniProt:Q56A08 |
| Proteins | Q8C5N3 | UniProt:Q8C5N3 |
| Proteins | Q8CEC6 | UniProt:Q8CEC6 |
| Proteins | Q9D6J3 | UniProt:Q9D6J3 |
| Proteins | Q9D198 | UniProt:Q9D198 |
| Proteins | Q9DBF7 | UniProt:Q9DBF7 |
| Proteins | Q9DBM1 | UniProt:Q9DBM1 |
| Proteins | A2ACQ1 | UniProt:A2ACQ1 |
| Proteins | Q3TWF6 | UniProt:Q3TWF6 |
| Proteins | Q9CXG3 | UniProt:Q9CXG3 |
| Proteins | Q5NCR9 | UniProt:Q5NCR9 |
| Proteins | Q9CS00 | UniProt:Q9CS00 |
| Proteins | Q91VN6 | UniProt:Q91VN6 |
| Proteins | Q8VDP2 | UniProt:Q8VDP2 |
| Proteins | A2A4P0 | UniProt:A2A4P0 |
| Proteins | Q8BM39 | UniProt:Q8BM39 |
| Proteins | Q9CWV6 | UniProt:Q9CWV6 |
| Proteins | Q9CR80 | UniProt:Q9CR80 |
| Proteins | Q8K1J5 | UniProt:Q8K1J5 |
| Proteins | Q9WV03 | UniProt:Q9WV03 |
| Proteins | Q9D0F4 | UniProt:Q9D0F4 |
| Proteins | Q3TQI7 | UniProt:Q3TQI7 |
| Proteins | Q9DB98 | UniProt:Q9DB98 |
| Proteins | Q64213 | UniProt:Q64213 |
| Proteins | Q8CH02 | UniProt:Q8CH02 |
| Proteins | Q5SUF2 | UniProt:Q5SUF2 |
| Proteins | Q8BGC0 | UniProt:Q8BGC0 |
| Proteins | Q9CQT2 | UniProt:Q9CQT2 Rbm7 |
| Proteins | Q9CZ96 | UniProt:Q9CZ96 |
| Proteins | Q3UZ01 | UniProt:Q3UZ01 |
| Proteins | Q9CQR5 | UniProt:Q9CQR5 |
| Proteins | Q9D361 | UniProt:Q9D361 |
| Proteins | Q9D384 | UniProt:Q9D384 |
| Proteins | Q8VIK1 | UniProt:Q8VIK1 |
| Proteins | Q9WTY1 | UniProt:Q9WTY1 |
| Proteins | Q62377 | UniProt:Q62377 |
